# The cytochrome P450 enzyme MpCYP78E1 inhibits meristem initiation and activity in *Marchantia polymorpha*

**DOI:** 10.1101/2025.04.28.651025

**Authors:** Victoria Spencer, Chloe Casey, Magdalena Mosiolek, Katharina Jandrasits, Natalie Edelbacher, Liam Dolan

**Affiliations:** Gregor Mendel Institute, Dr-Bohr-Gasse 3, 1030 Vienna, Austria; St Anna Children’s Cancer Research Institute (CCRI), Zimmermannplatz 10, 1090 Vienna, Austria

**Keywords:** Shoot branching, meristem, dichotomous branching, bryophytes, liverworts, Marchantia polymorpha, cytochrome P450, gametangiophores

## Abstract

Plant shoot branches are formed by the initiation and activation of generative centres known as meristems. In dichotomously branching plants, such as many bryophytes and lycophytes, new meristems are formed when a pre-existing meristem splits into two daughter meristems. These meristems may be active and produce shoot branches or may be inactive. Here, we show that in conditions where meristem inactivation occurs, such as simulated shade, the position of the inactive meristem alternated between either side of the plant body in the liverwort *Marchantia polymorpha.* Using this predictable pattern, we generated transcriptomes of active and inactive meristems and identified the cytochrome P450 monooxygenase, MpCYP78E1, as a novel regulator of meristem activity. Mp*CYP78E1* reporter expression was higher in active meristems than inactive meristems. More meristems were active in loss of function mutants than wild type, and fewer meristems were active in gain of function mutants, indicating that MpCYP78E1 inhibits meristem activity. Furthermore, unlike wild type, Mp*cyp78e1* loss of function mutants produced supernumerary meristem from the centre of the mature plant body. We conclude that MpCYP78E1 inhibits both meristem initiation and activity to modulate shoot branching architecture.

## INTRODUCTION

The architecture of plants is generated by the initiation and activity of meristems; generative structures comprising stem cells and proliferative cells that differentiate to produce new tissues and organs. During shoot branching, a new meristem is formed, which depending on endogenous and environmental signals may be active and form a new growth axis or may remain inactive (Li *et al*., 2024). Two distinct modes of shoot branching exist in land plants. Seed plants branch sub-apically in a process known as axillary branching; new meristems initiate behind the apex in the axils of developing leaves (Domagalska and Leyser, 2011; Luo *et al*., 2021). Other plant lineages such as many bryophytes, lycophytes and ferns branch apically in a process known as dichotomous branching; new branches are produced directly at the shoot apex through the splitting of the meristem to form two daughter meristems (Gola, 2014; Harrison, 2017; Spencer *et al*., 2021). Dichotomous branching was likely ancestral in land plants and axillary branching evolved independently numerous times (Chomicki *et al*., 2017; Harrison and Morris, 2018). The molecular mechanisms controlling dichotomous branching are poorly understood.

Dichotomous branching occurs in the haploid stage of the lifecycle in the liverwort, *Marchantia polymorpha*, from meristems located within notches at the apices of the thalloid plant body (Solly *et al*., 2017; Hirakawa *et al*., 2020; Spencer *et al*., 2024). The activity of meristems is modulated by environmental conditions such as light and shade. For example, when *M. polymorpha* gemmae are grown in white light, the apical meristem splits to produce two new daughter meristems, which both remain active and each produce a shoot branch (Streubel *et al*., 2023). Branching is therefore isotomous (equal) early in the development of the plant (Harrison and Morris, 2018; Streubel *et al*., 2023). After approximately six cycles of dichotomy, the two new daughter meristems develop different fates – one meristem becomes active and the other becomes inactive (or dormant) – likely to avoid the effects of self-shading (Streubel *et al*., 2023). Branching is therefore anisotomous (unequal). When plants are grown in simulated shade, inactive meristems develop earlier (before the sixth cycle of dichotomy). Meristem inactivation requires the *SQUAMOSA PROMOTER BINDING PROTEIN-LIKE 1* (Mp*SPL1*) transcription factor (Streubel *et al*., 2023), and is maintained by apically produced auxin that moves through the plant to inhibit neighbouring meristems (Binns and Maravolo, 1972; Davidonis and Munroe, 1972; Maravolo, 1976; Streubel *et al*., 2023). Later in development, the apical meristem undergoes a transition to form a determinate gamete-producing branch known as a gametangiophore. Meristem activity in the developing thallus is therefore modulated by a range of factors including plant age and environment.

Here, we report the discovery of a novel repressor of meristem initiation and activity. We identified the cytochrome P450 enzyme, MpCYP78E1, by comparing the transcriptomes of active and inactive meristems. Mp*CYP78E1* mRNA levels are higher in active meristems than inactive meristems, and MpCYP78E1 inhibits meristem activity in both white light and simulated shade. Furthermore, MpCYP78E1 inhibits the initiation of new meristems from non-meristematic tissue at the centre of the mature plant body. We conclude that MpCYP78E1 represses meristem initiation and meristem activity and is a key regulator of shoot branching architecture in *M. polymorpha*.

## RESULTS

### Active and inactive meristems alternate positions along the thallus axis

*M. polymorpha* plants comprise thallus axes derived from apical meristems. Apical meristems duplicate during dichotomous branching and each resulting meristem may be active or inactive. The proportion of inactive meristems in a plant is higher in simulated shade than in white light (Streubel *et al*., 2023). Therefore, we used simulated shade experimental conditions to discover regulators of meristem activity. Active meristems are located at the thallus apex and produce vegetative thallus branches. By contrast, inactive meristems do not develop new tissue and are positioned below the apex to the side of the thallus body. The distance between the active meristem and the inactive meristem increases as the active meristem produces new tissue and the thallus between these meristems elongates.

To determine the spatial pattern of active and inactive meristems along the thallus, we quantified the number and position of active and inactive meristems in 8-week old wild type Tak-1 plants (Fig. 1). Plants were grown in white light for 2 weeks followed by simulated shade for 6 weeks and were grown from gemmae – disc-shaped vegetative propagules with two meristems, one on either side of the disc. After 2 weeks in white light, an average of 8 active meristems (8.3 ± 1.3, n=60) and no inactive meristems were observed (Fig. 1B-C’, G). When plants were grown for 4 weeks in white light, all meristems were active (29.7 ± 7.2, n=4; Fig. S1A). In comparison, after 2 weeks in white light followed by 2 weeks in simulated shade (total 4 weeks), 84 % of meristems were active (20.8 ± 3.6, n=60) and 14 % of meristems were inactive (3.3 ± 1.9, n=60) (Fig. 1B, D, D’). After 2 weeks in white light and 4 weeks in simulated shade (total 6 weeks), 41 % of meristems were active (20.9 ± 3.8, n=60) and 23 % of the meristems were inactive (12.1 ± 6.4, n=60). This is consistent with the observation that meristem inactivation is induced by simulated shade (Streubel *et al*., 2023). Furthermore, we observed that if the meristem on the left side of the thallus was inactive following a dichotomy event, the meristem on the right became inactive after the next dichotomy event (Fig. 1D-E’). The position of the inactive meristem therefore alternated from the left to right side of the thallus axis as development proceeded. The repetition of these anisotomous (unequal) branching events generated a pseudomonopodial axis (Fig. 1H). We conclude that the position of active and inactive meristems is predictable.

**Figure 1:**
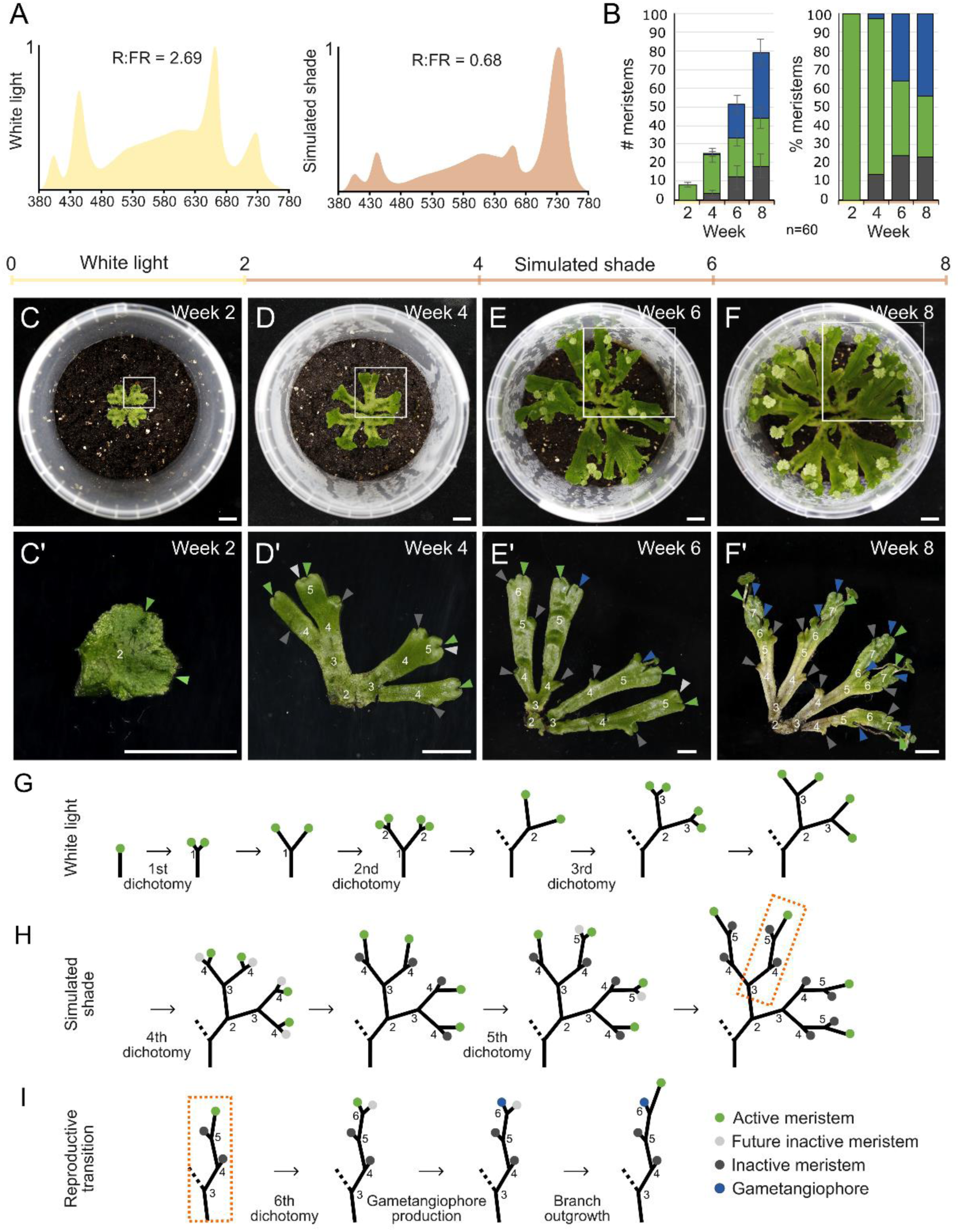
The positions of active and inactive meristems alternate along the *Marchantia polymorpha* thallus body. (**A**) Typical white light and simulated shade spectra. In white light, the red: far-red ratio and light intensity are high. In simulated shade, the red: far-red ratio and light intensity are low. (**B**) Wild type Tak-1 was grown for 2 weeks in white light followed by 6 weeks in simulated shade. The numbers of active meristems (green), inactive meristems (dark grey), and reproductive structures (gametangiophores) (blue) were quantified. Gametangiophores were only produced after transfer to shade. Notches were counted as a proxy for meristems. The experiment was repeated twice and showed consistent results. The data presented are pooled from the two experiments. n= 60. (**C**-**F’**) Images of plants at week 2 (C, C’), week 4 (D, D’), week 6 (E, E’) and week 8 (F, F’). C, D, E and F show the whole thallus and C’, D’, E’ and F’ show the branching architecture of one thallus quarter. (**C**, **C’**, **G**) In white light, *M. polymorpha* wild type Tak-1 produced an isotomously branching plant body, in which the apical meristem produced two daughter meristems by dichotomy, which both remained active (green circles). Each meristem produced a branch, which grew equally. (**D**, **D’**, **H**) After transfer to shade, plants produced an anisotomously branching plant body. One of the daughter meristems remained active (green circles) whilst the other became inactive (dark grey circles). The position of the inactive meristem alternated from the left to the right, and therefore the “future inactive meristem” (light grey circles) could be predicted before it displayed any morphological difference to the active meristem. (**E**, **E’**, **F**, **F’**, **I**) Based on the alternation of active and inactive meristem, gametangiophores are predicted to form from the active meristem. Black lines= branching axes. Scale bars = 10 mm.

After prolonged growth in simulated shade, thallus apical meristems of wild type Tak-1 and Tak-2 plants underwent the reproductive transition and produced gametangiophore meristems. Gametangiophores are modified, determinate thalli that develop egg-producing archegonia or sperm-producing antheridia in females and males respectively (Shimamura, 2016). Gametangiophore initiation was observed at week 6 and each developed from a single meristem produced by meristem dichotomy at the thallus apex (Fig. 1B, E-F’, Fig. S1B). In these conditions, the gametangiophore meristems were produced from the predicted active meristem after seven dichotomy events (6.7 ± 1.4, n=10) (Fig. S1B-D). The development of a gametangiophore meristem from an active meristem was accompanied by the activation of the neighbouring meristem which would otherwise have been inactive (Fig 1F, F’, I). We conclude that the gametangiophore meristem develops from the active meristem following dichotomy.

### Mp*CYP78E1* mRNA levels are higher in active meristems than inactive meristems

To identify genes that regulate meristem activity, we generated transcriptomes from pairs of neighbouring active and inactive meristems from wild type Tak-1 plants at week 4 (Fig. 2). Notches containing the active meristem (active notch) and notches containing the inactive meristem (inactive notch) were harvested at three stages following dichotomy. At stage 1, the active notch and the inactive notch were located next to each other and were morphologically indistinguishable (Fig. 2A). At stage 2, the inactive notch was positioned on the side of the thallus less than 5 mm from the active notch. At stage 3, the inactive notch was more than 5 mm from the active notch.

**Figure 2:**
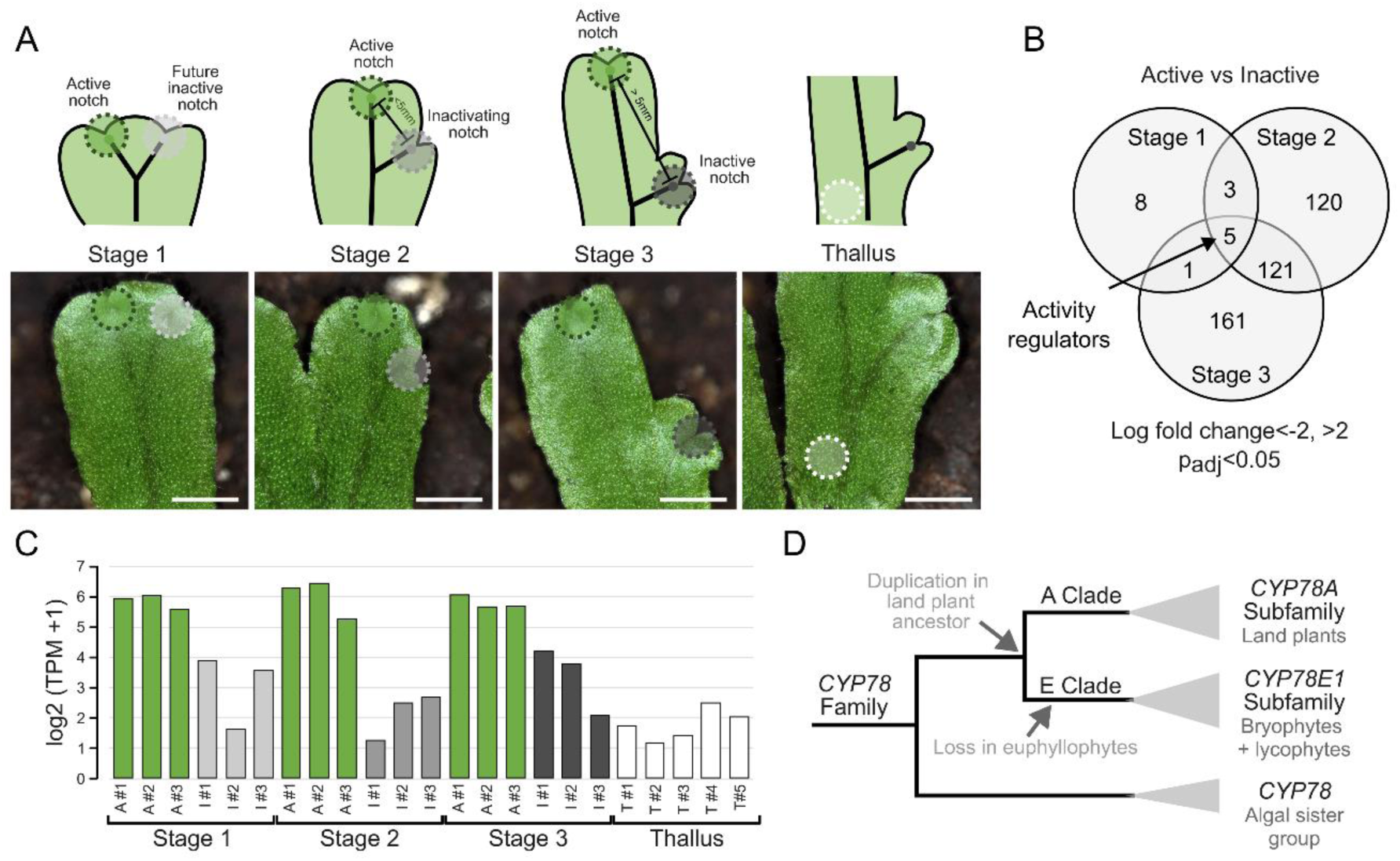
*MpCYP78E1* was identified as a candidate regulator of meristem activity by RNA-Seq. (**A**) The active notch (containing an active meristem) and the inactive notch (containing an inactive meristem) of wild type Tak-1 were harvested at three stages of meristem inactivation. Stage 1 was recently after a dichotomy event, in which the active and future inactive notch were < 3mm apart. Stage 2 was later after dichotomy, in which the inactivating notch was positioned on the side of the thallus body, within 3-5 mm of the active notch. At stage 3, the inactive notch was more than 5 mm from the active notch. Thallus tissue was harvested as a non-meristem control. Scale bar= 2 mm. (**B**) Venn diagram showing the Differentially Expressed Genes (DEGs) between active and inactive notches at the three stages of inactivation. 5 genes were differentially expressed in all three stages and were identified as potential meristem activity regulators. (**C**) Log2 (TPM +1) expression of Mp1g14150 (Mp*CYP78E1*) in each sample. n= 3 for active (A) and inactive (I) notches, and n= 5 for thallus (T). (**D**) Summary cladogram of CYTOCHROME P450 family 78 based on phylogeny generated by SHOOT.bio (Fig. S3). Mp*CYP78E1* is within the CYP78 E subfamily, which contains only non-seed plant lineages. The closest sister group is the CYP78 A subfamily, which includes an *M. polymorpha* ortholog (Mp3g23930), as well as *Arabidopsis thaliana* genes such as At*KLUH*.

Genes that were differentially expressed (DEGs) between active and inactive meristems in paired notches (Log2 Fold Change > 2 or < -2 and a p_adjusted_< 0.05) were identified (Fig. 2B). 17, 249, and 288 DEGs were found for stages 1, 2 and 3 respectively. 5 genes were differentially expressed in all three stages of meristem inactivation (Mp4g05770; Mp3g25440: MLP-LIKE PROTEIN 423; Mp2g18830: cytolysin/lectin; Mp1g14150: cytochrome P450 family 78 subfamily E; Mp1g28130: O-methyltransferase; Fig. S2). Each of these genes were expressed more strongly in notches containing active meristems than notches containing inactive meristems (Fig. 2C, Fig. S2A). Of these 5 genes, we selected Mp3g25440 and Mp1g14150 because independent data in the MarpolBase expression database verified that the corresponding transcripts were enriched in the meristem region of the thallus (Fig. S2B). We generated loss of function mutant lines for Mp3g25440 and Mp1g14150 using CRISPR/Cas9. Since the branching and thallus morphology of Mp3g25440 was similar to wild type (Fig. S3), we proceeded only with Mp1g14150 for further analysis.

Mp1g14150 is a member of the CYTOCHROME P450 superfamily of monooxygenases that are present in all eukaryotes and oxidise diverse substrates in numerous metabolic pathways. Mp1g14150 belongs to CYTOCHROME P450 clan 71, family 78 and subfamily E and was therefore named Mp*CYP78E1* according to standard conventions (Casey and Dolan, 2023). A gene tree generated using SHOOT (Emms and Kelly, 2022) confirmed this classification and found that the CYP78E subfamily clade comprised only Mp*CYP78E1* and other bryophyte and lycophyte orthologues (Fig. 2D, Fig. S4). No fern or seed plant genes were identified within this subfamily, and the most closely related euphyllophyte genes were members of the paralogous CYP78A clade. We therefore hypothesise that the CYP78E subfamily was present in the common ancestor of land plants and was lost after the divergence of lycophytes from the lineage that gave rise to the ferns and seed plants.

### Mp*CYP78E1* expression is enriched in active meristems

To identify where Mp*CYP78E1* is expressed within active and inactive meristems, we first described the morphology and anatomy of active and inactive notches of wild type Tak-1 plants at week 4 (2 weeks white light + 2 weeks simulated shade) (Fig. 3). In 38 % of the active notches, two neighbouring meristems were present, because of a recent dichotomy event (Fig. 3A, E). 85 % of active notches contained gametangiophore meristems at week 4 (Fig. 3A, F). Inactive notches always contained only a single active meristem and no gametangiophore meristem. Active and inactive meristems exhibited the same U-shaped structure, with the stem cell niche positioned at the base of the notch in the outer cell layer (Fig. 3A-D).

**Figure 3:**
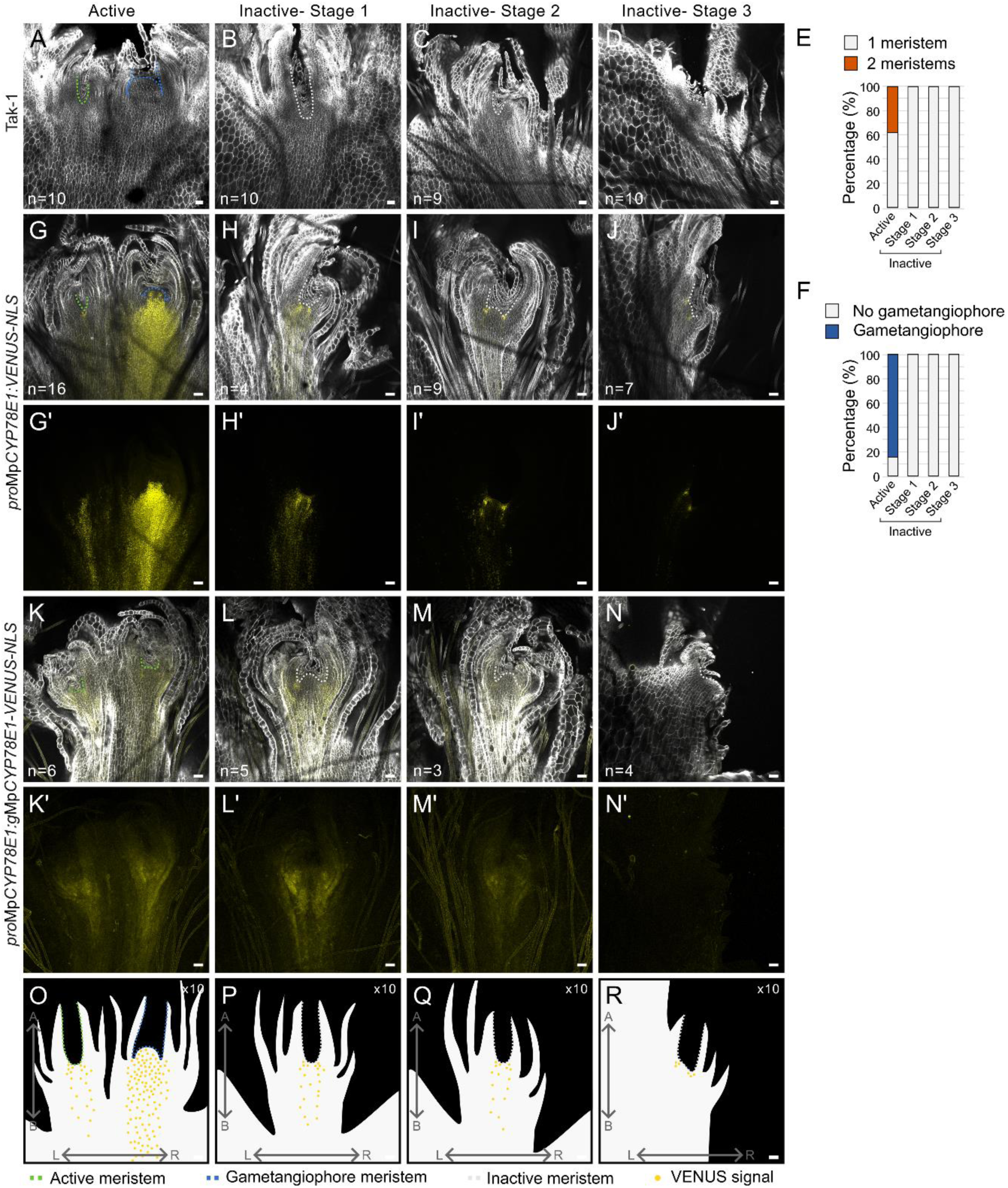
Mp*CYP78E1* expression is enriched in active meristems and gametangiophore meristems. (**A-D**) Confocal images of harvested wild type Tak-1 meristems in active, inactive stage 1, inactive stage 2 and inactive stage 3 notches. Samples were fixed and cleared before imaging. (**E**) Quantification of the number of meristems within each stage. Only the active meristem underwent dichotomy. (**F**) Quantification of the number of samples producing gametangiophore meristems. No gametangiophores were evident when harvesting, however they are evident when fixed, cleared and imaged. Only the active notch produced gametangiophore meristems, consistent with the observations described in Fig. 1. (**G**-**J’’**) Confocal images of fixed and cleared meristems of *pro*Mp*CYP78E1:VENUS-NLS* lines at different stages of meristem inactivation. G, H, I, and J show optical sections in the frontal plane. G’, H’, I’ and J’ show z-projections of the reporter signal across a Z-stack of the meristem. There is strong signal accumulation around each meristem and this extended basally from the meristem in active (green), stage 1 inactive and stage 2 inactive (grey) meristems. Signal was enriched in the gametangiophore meristems (blue). The experiment was repeated twice and showed consistent results. Sample number is pooled from both replicates and is shown in the figure. A second independent line is shown in Fig. S5. (**K**-**N’’**) Confocal images of fixed and cleared meristems of *pro*Mp*CYP78E1:gMpCYP78E1-VENUS* lines at different stages of meristem inactivation. K, L, M, and N show optical sections in the frontal plane. K’, L’, M’ and N’ show z-projections of the reporter signal across a Z-stack of the meristem. There was strong signal accumulation around all meristems and this extended basally towards the midrib in all samples except the stage 3 inactive meristems. A second and third independent line are shown in Fig. S5. (**O**-**R**) O, P, Q and R show schematics of typical reporter expression. Cell wall = white. VENUS signal = yellow. Scale bars = 50 mm.

To independently verify that Mp*CYP78E1* transcripts were enriched in active notches, we generated transcriptional reporter lines using the 5 kb region upstream (5’) of the start codon to drive the transcription of NLS-tagged *VENUS*. Consistent expression patterns were observed in 12 independently transformed lines. Two lines (*pro*Mp*CYP78E1:VENUS-NLS^1m^*and *pro*Mp*CYP78E1:VENUS-NLS^8f^*) were selected for further analysis and notches were collected at week 4 (Fig. 3G-J’, Fig. S5A-D’). VENUS signal was enriched in the active notches, both in the active meristems and the gametangiophore meristems. Signal was also present in stage 1 and stage 2 inactive meristems, but by stage 3 expression was weaker and was restricted to a small region around the apical cells of the inactive meristem. These data indicate that MpCYP78E1 transcripts are expressed at high levels in active meristems and low levels in stage 3 inactive meristems.

To define the cells where MpCYP78E1 protein accumulates, a *pro*Mp*CYP78E1:g*Mp*CYP78E1-VENUS* construct was transformed into the Mp*cyp78e1^LOF_g2_14m^* mutant background. Three independent complementation lines (Mp*cyp78e1^COMP^*) were obtained, and their VENUS localisation was analysed at week 4 (Fig. 3K-N’, Fig. S5E-L’). VENUS signal accumulated around the stem cell niche of all meristems, and extended basally towards the midrib in active, stage 1 inactive, and stage 2 inactive meristems, but not stage 3 inactive meristems. These results are consistent with the transcriptional reporter lines (Fig. 3 G-J’), supporting the conclusion that MpCYP78E1 is enriched in active meristems, but becomes progressively restricted in the inactive meristem.

### MpCYP78E1 inhibits meristem activity

To test the hypothesis that Mp*CYP78E1* regulates meristem activity in *M. polymorpha*, CRISPR Mp*cyp78e1^LOSS^ ^OF^ ^FUNCTION^ ^(LOF)^* mutants were generated (Fig. 4). gRNAs were designed targeting the membrane associated domain encoded in the first exon and the CYTOCHROME P450 domain encoded in the second exon. From each gRNA transformation, 3 independent lines were selected for further analysis (Fig. S6). To investigate if MpCYP78E1 regulates meristem activity, wild type and Mp*cyp78e1^LOF^* mutant plants were grown and active meristems, inactive meristems and gametangiophores were counted at weeks 4 and 8 (Fig. 4A-D). In Mp*cyp78e1^LOF^* mutants at week 4, 2-10 % of meristems were inactive, compared to 25-34 % of wild type meristems (Fig. 5A, C), indicating that *MpCYP78E1* promotes meristem inactivation. Since only active meristems produce gametangiophores (Fig. 1, 3) and more meristems were active in the Mp*cyp78e1^LOF^* mutants than wild type, we predicted that the Mp*cyp78e1^LOF^* mutants would produce more gametangiophores than wild type. Consistent with this hypothesis, we found that 89-98 % of meristems developed gametangiophores in the Mp*cyp78e1^LOF^* mutant lines by week 8, compared to 29-36 % of meristems in wild type (Fig. 4B, D). These data support the hypothesis that MpCYP78E1 negatively regulates meristem activity.

**Figure 4:**
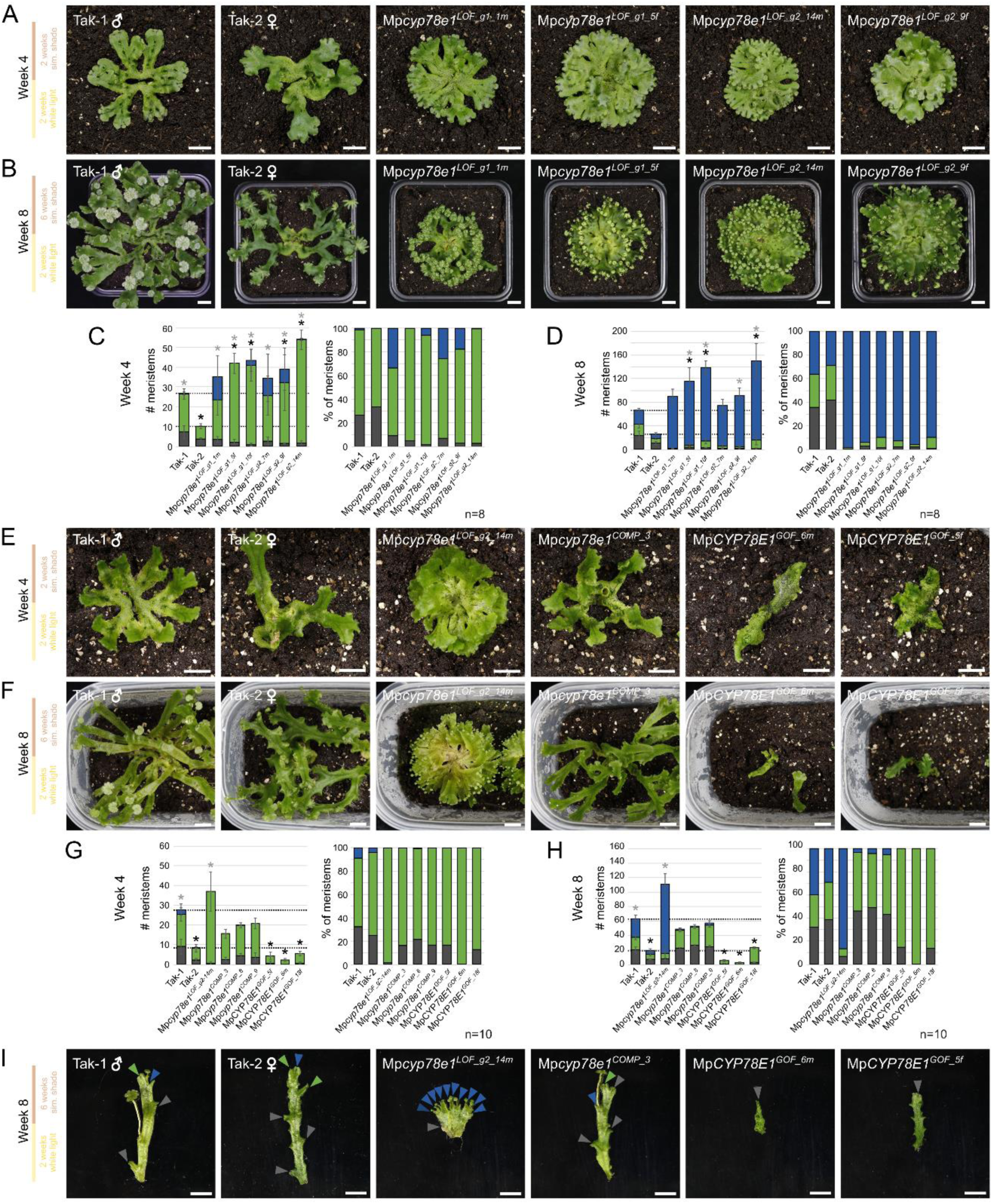
MpCYP78E1 inhibits meristem activity in simulated shade. (**A**) Wild type Tak-1, wild type Tak-2, and 6 independent Mp*cyp78e1^LOF^* lines were grown for two weeks in white light and two weeks in simulated shade (4 weeks total). Inactive meristems were formed in wild type plants, but rarely in Mp*cyp78e1^LOF^* lines. (**B**) After two weeks in white light and six weeks in simulated shade (8 weeks total), wild type and mutant plants both made reproductive structures. (**C**-**D**) The total number and percentage of meristems that were active (green), inactive (dark grey), or reproductive (blue) was quantified at week 4 (C) and week 8 (D). At week 4, ∼25-34 % of wild type Tak-1 and Tak-2 meristems were inactive, and less than 10 % of meristems were inactive in all six Mp*cyp78e1^LOF^* lines. At week 8, 26-36 % of wild type meristems were reproductive, whilst this was greater than 89 % in all six Mp*cyp78e1^LOF^*lines. A one-way ANOVA with Tukey’s HSD multiple comparison test was performed for the total number of meristems at week 4 (F(7,56)=37.338, p=7.32E-19) and a Kruskal-Wallis with Dunn’s multiple comparison test was performed for the total number of meristems at week 8 [χ^2^(7)=54.3, p=2.06E-9]. n= 8 and error bars indicate standard deviation. Black * denotes a significant difference to Tak-1 (p-value <0.05) and grey * denotes a significant difference to Tak-2 (p-value <0.05). (**E**) Wild type Tak-1, Tak-2, 3 independent complementation lines and 3 independent Mp*CYP78E1^GOF^* lines were grown for two weeks in white light and two weeks in simulated shade (4 weeks total). Inactive meristems were formed in wild type plants, complementation lines, and Mp*CYP78E1^GOF^* lines. (**F**) After two weeks in white light and six weeks in simulated shade (8 weeks total), wild type and complementation lines made reproductive structures, whilst Mp*CYP78E1^GOF^* lines did not. (**G-H**) The total number and percentage of meristems that were active, inactive, or reproductive was quantified at week 4 (G) and week 8 (H). Complementation of the Mp*cyp78e1^LOF^* mutants with the translational reporter construct restored the number and percentage of each meristem type to intermediate of Tak-1 and Tak-2. The total number of meristems was lower in Mp*CYP78E1^GOF^* lines than wild type. 29-41 % of meristems in wild type Tak-1 and Tak-2 lines were reproductive, 87 % were reproductive in Mp*cyp78e1^LOF_g2_14m^*, whilst 0 % of meristems produced reproductive structures in Mp*CYP78E1^GOF^* lines. The experiment was repeated twice and showed consistent results. The data presented are pooled from the two experiments. Kruskal-Wallis with Dunn’s multiple comparison tests were performed for the total number of meristems at week 4 and week 8 respectively [χ^2^(8)=79.4, p=6.34E-14; χ^2^(8)=82, p=1.93E-14]. n= 10 in total and error bars indicate standard deviation. Black * denotes a significant difference to Tak-1 (p-value <0.05) and grey * denotes a significant difference to Tak-2 (p-value <0.05). (**I**) Branches from the 3 plastochron were dissected from plants grown for 8 weeks. Scale bars = 10 mm.

**Figure 5:**
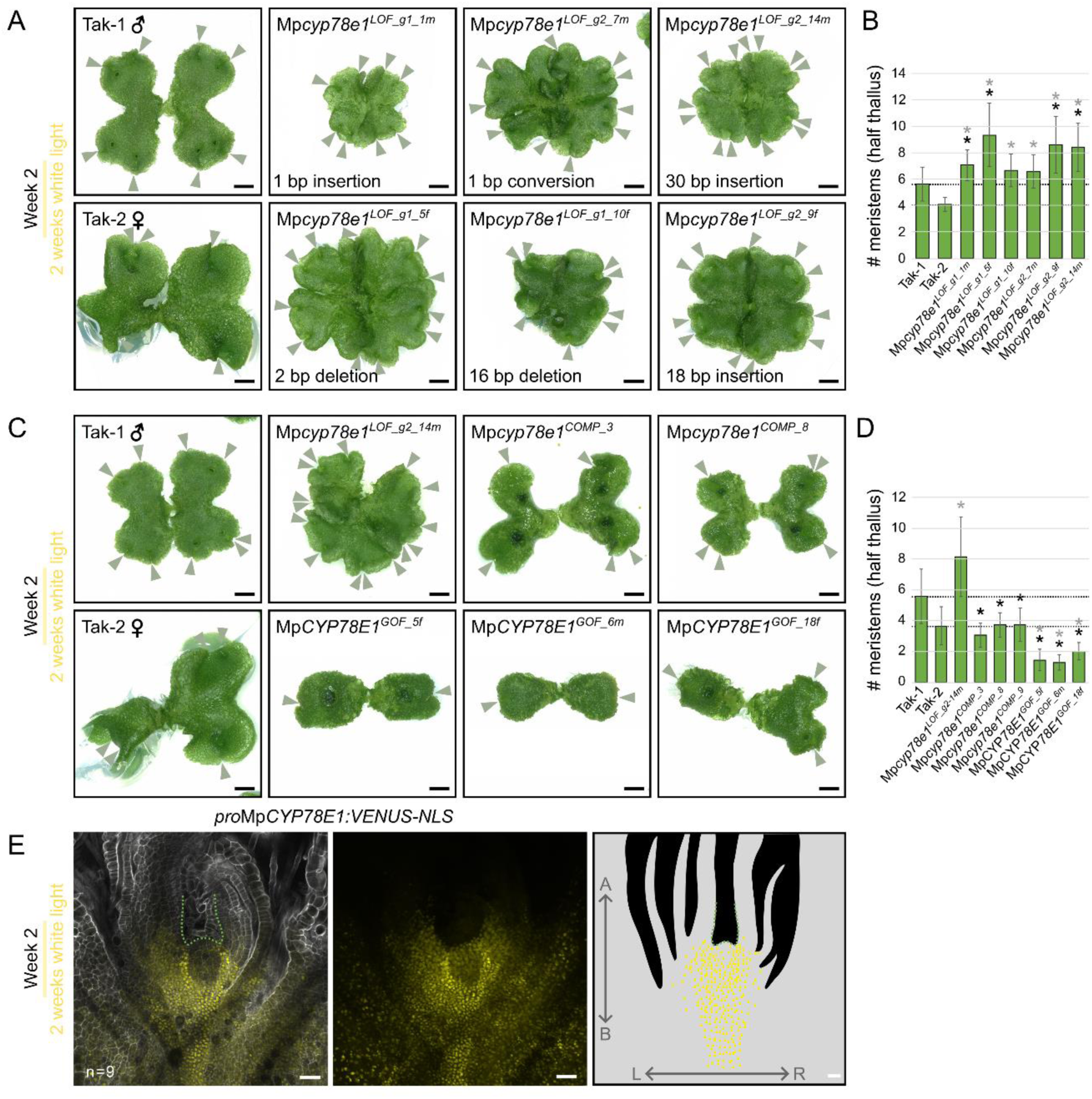
MpCYP78E1 inhibits meristem initiation in white light. (**A**) Wild type Tak-1 and Tak-2, and 6 independent Mp*cyp78e1^LOF^* lines were grown for two weeks in white light. (**B**) Meristem number was counted at week 2. There were more meristems in all 6 mutant lines than Tak-1 and Tak-2. The experiment was repeated twice and showed consistent results. The data presented are pooled from the two experiments. A Kruskal-Wallis with Dunn’s multiple comparison test was performed for the total number of meristems [χ^2^(7)=103.0, p=2.34E-19]. n= 34, 18, 23, 26, 26, 24, 21 and 26 respectively and error bars indicate standard deviation. Black * denotes a significant difference to Tak-1 (p-value <0.05) and grey * denotes a significant difference to Tak-2 (p-value <0.05). (**C**) Wild type Tak-1 and Tak-2, a Mp*cyp78e1^LOF^* line, 3 complementation lines, and 3 Mp*CYP78E1^GOF^*line were grown for two weeks in white light. (**D**) Meristem number was counted at week 2. There were fewer meristems in all 3 GOF lines than both Tak-1 and Tak-2. The experiment was repeated twice and showed consistent results. The data presented are pooled from the two experiments. A Kruskal-Wallis with Dunn’s multiple comparison test was performed for the total number of meristems [χ^2^(8)=273, p=2.68E-54]. n= 45, 32, 27, 47, 45, 42, 42, 45 and 41 respectively and error bars indicate standard deviation. Black * denotes a significant difference to Tak-1 (p-value <0.05) and grey * denotes a significant difference to Tak-2 (p-value <0.05). Scale bars for A and C = 2.5 mm. (**E**) Confocal image of a fixed and cleared meristem of a *pro*Mp*CYP78E1:VENUS-NLS* line grown for 2 weeks in white light. Left, an optical section in the frontal plane; middle, a z-projection of the reporter signal across a Z-stack of the meristem; and right, a schematic of typical reporter expression. There is strong signal accumulation around each stem cell niche and extending back from the meristem. Cell wall= white. *pro*Mp*CYP78E1:VENUS-NLS* signal= yellow. A second independent line is shown in Fig. S8. Scale bars = 50 µm.

To verify that the mutations in *MpCYP78E1* cause the defective meristem phenotypes, we characterised the phenotype of 3 independent lines expressing a complementing gRNA-insensitive *pro*Mp*CYP78E1:g*Mp*CYP78E1-VENUS* construct in the Mp*cyp78e1^LOF^* background at week 4 and 8 (Fig. 4E-H; expression shown in Fig. 3). At both time points the total number of meristems in the complementation lines was within the range of wild type Tak-1 and Tak-2 (Fig. 4E, G), as would be expected if the wild type phenotype was restored by the transgene. 14-22 % of meristems were inactive in Mp*cyp78e1^COMP^* lines, which was more similar to wild type (26-32 %) than the Mp*cyp78e1^LOF^*mutant (2 %) at week 4. The Mp*CYP78E1* transgene therefore restores the wild type phenotype and confirms that the Mp*cyp78e1^LOF^* mutation causes the defective meristem phenotype. This further supports the conclusion that MpCYP78E1 negatively regulates meristem activity.

If MpCYP78E1 represses meristem activity, we predicted that meristem activity would be reduced in plants overexpressing Mp*CYP78E1* (Mp*CYP78E1^GAIN^ ^OF^ ^FUNCTION^ ^(GOF)^*) compared to wild type. Inactive meristems do not undergo dichotomy, so we would expect that overexpression of Mp*CYP78E1* would reduce the number of dichotomy events and therefore reduce the total number of meristems in the whole plant. To test this hypothesis, we expressed the *pro*Mp*CYP78E1:g*Mp*CYP78E1-VENUS* construct in the wild type spores produced from a cross between Tak-1 x Tak-2 and generated over 30 independent lines. 3 lines with strong *VENUS* expression and consistent phenotypes were selected for further analysis (Fig. S7). The numbers of active and inactive meristems were quantified in the wild type and Mp*CYP78E1^GOF^* lines at week 4 and 8 (Fig. 4E-H). There were fewer meristems in Mp*CYP78E1^GOF^* lines than wild type. Furthermore, in wild type, only active meristems produce reproductive structures after sustained exposure to simulated shade (Fig. 1, 3). If meristem activity was reduced in the meristems of Mp*CYP78E1^GOF^* lines, we would expect fewer meristems to produce gametangiophores than wild type. Consistently, no reproductive structures were formed in any of the Mp*CYP78E1^GOF^* lines after 8 weeks of growth (Fig. 4F, H). We therefore conclude that Mp*CYP78E1^GOF^* apical meristems were mostly inactive. Taken together, the phenotypes of Mp*cyp78e1^LOF^* and Mp*CYP78E1^GOF^* lines support the conclusion that MpCYP78E1 is an inhibitor of meristem activity.

### MpCYP78E1 is a general inhibitor of meristem activity

We found that MpCYP78E1 inhibits meristem activity in plants grown in simulated shade. If MpCYP78E1 activity is only induced under these conditions, then both Mp*cyp78e1^LOF^* and Mp*CYP78E1^GOF^*mutants should develop the same number of meristems as wild type when grown in white light. Mutants were grown for 2 weeks in white light (Fig. 5). The total number of active meristems was higher in Mp*cyp78e1^LOF^* lines than wild type and lower in Mp*CYP78E1^GOF^* lines than wild type (Fig. 5A-D). This demonstrates that MpCYP78E1 inhibits meristem activity in white light as well as simulated shade. Consistent with this, *pro*Mp*CYP78E1:VENUS-NLS* lines grown for two weeks in white light accumulated VENUS signal in active meristems (Fig. 5E, Fig. S8). We conclude that MpCYP78E1 is a general inhibitor of meristem activity and not a shade specific repressor of meristem activity.

### MpCYP78E1 represses meristem initiation

Meristems are typically initiated during dichotomy of the active meristem in *M. polymorpha*, but also initiate from non-meristematic tissues following damage or removal of the active meristem (Binns and Maravolo, 1972; Streubel *et al*., 2023). Since MpCYP78E1 inhibits meristem initiation during dichotomy, we speculated that MpCYP78E1 may be a general inhibitor of meristem initiation. In plants grown for 3 weeks in white light, we observed that new branches (and associated meristems) initiated from the central of the thallus in 0 % and 29 % of wild type Tak-1 and Tak-2 plants, respectively (Fig. 6). Fewer plants produced supernumerary branches in Mp*CYP78E1^GOF^* lines than wild type (0-17 %), and more plants produced supernumerary branches in Mp*cyp78e1^LOF^* mutant lines than wild type (67-81 %) (Fig. 6). These data are consistent with the hypothesis that MpCYP78E1 inhibits meristem initiation from non-meristematic tissues. MpCYP78E1 is therefore a general inhibitor of meristem initiation.

**Figure 6:**
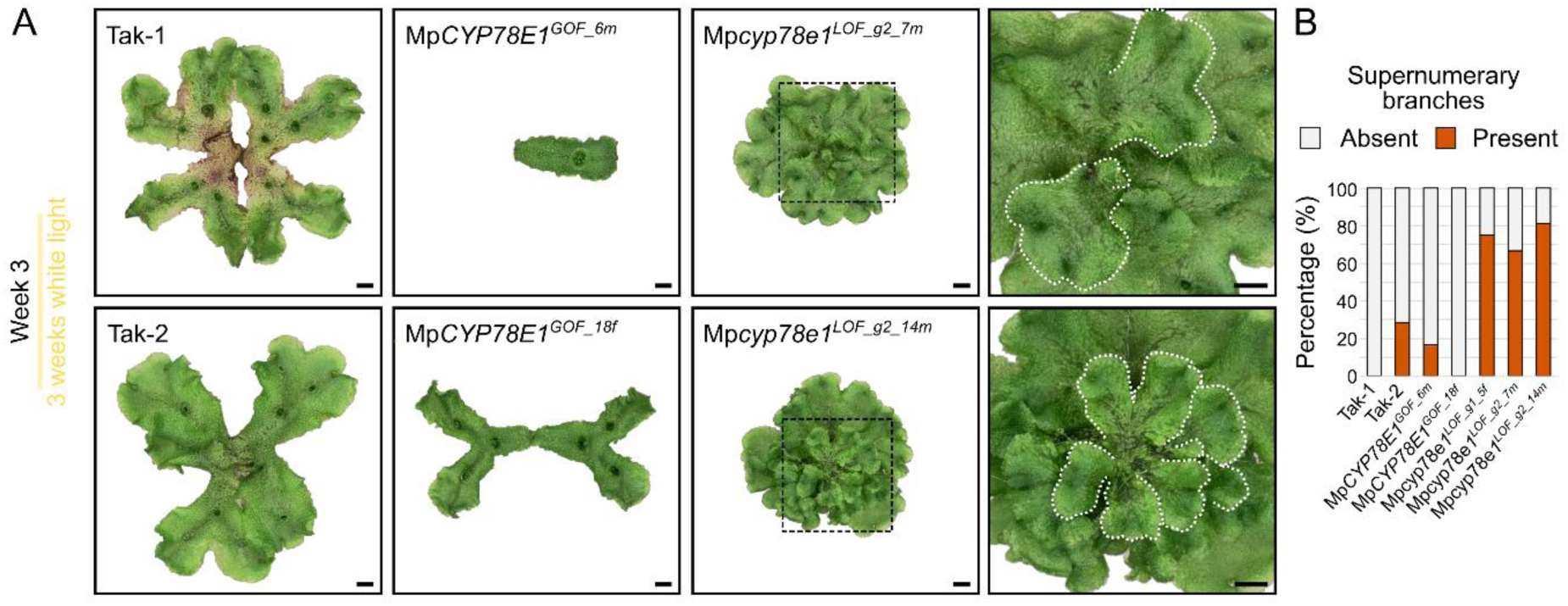
MpCYP78E1 inhibits meristem initiation in white light. (**A**) Photographs of wild type Tak-1, Tak-2, Mp*CYP78E1^GOF^*lines and Mp*cyp78e1^LOF^* lines after three weeks of growth in white light. Supernumerary thallus branches (white dashed line) were produced from the central plant body of Mp*cyp78e1^LOF^* lines. Black dotted box is magnified to the right. The background was removed for clarity and the original images are shown in Fig. S9. Scale bars = 2.5 mm. (**B**) The percentage of plants with supernumerary branches. Wild type and gain of function lines rarely produced supernumerary branches (0-29 %), whilst all three independent loss of function lines produced supernumerary branches in most plants (67-81 %). n= 16, 14, 12, 14, 8, 15 and 16 respectively.

## DISCUSSION

The branching architecture of plants depends on the pattern of meristem initiation and the activity of these meristems once formed (Li *et al*., 2024). The axes of the *M. polymorpha* thallus produce new meristems at their apex through dichotomy (Solly *et al*., 2017; Hirakawa *et al*., 2020). A single meristem expands, splits, and the resulting two daughter meristems may be active or inactive (Davidonis and Munroe, 1972; Streubel *et al*., 2023). We discovered that MpCYP78E1 (Mp1g14150) inhibits meristem initiation; there is an increased rate of dichotomy in active meristems of Mp*cyp78e1* loss of function mutants compared to wild type, and loss of function mutants also produce supernumerary meristems from non-meristematic tissues at the centre of the mature plant. Consistently, plants overexpressing Mp*CYP78E1* produce fewer total meristems than wild type. Furthermore, MpCYP78E1 represses the activity of meristems; almost all meristems are active in Mp*cyp78e1* loss of function mutants grown in simulated shade, unlike wild type which produces active and inactive meristems. Since Mp*CYP78E1* expression is enriched in active meristems and MpCYP78E1 inhibits both meristem initiation and activity, we speculate that MpCYP78E1 in active meristems produces a mobile signal which inhibits the initiation and activity of neighbouring meristems.

MpCYP78E1 is a member of the Cytochrome P450 superfamily of enzymes, which are present in all kingdoms of life (Chapple, 1998; Fang *et al*., 2024). These enzymes are heme-containing proteins that most commonly catalyse oxidation reactions of organic substrates. In plants, they are involved in the biosynthesis of diverse classes of molecules including hormones, secondary metabolites, and pigments, as well as the metabolism of molecules such as fatty acids (Hansen *et al*., 2021). Mp*CYP78E1* is a member of the plant-specific CYP78 family, which comprises subclade E (containing Mp*CYP78E1*) and subclade A (Casey and Dolan, 2023). Phylogenetic analysis identified subclade E genes in the bryophytes and lycophytes but not in the euphyllophytes. Based on this analysis, we speculate that this gene was present in the common ancestor of land plants but was lost after the divergence of lycophytes from ferns and seed plants. This hypothesised loss coincides with reduced complexity of dichotomously branching life-stages in the euphyllophyte lineage (Chomicki *et al*., 2017; Harrison and Morris, 2018). There are no reports of the function of E subclade members in the CYP78 family. However, genes of the paralogous CYP78 A subclade have been characterised. The subclade A members, *KLUH/ CYP78A5* and *CYP78A7* in *Arabidopsis thaliana* and their ortholog *PLASTOCHRON1* (*PLA1*) in *Oryza sativa,* regulate organ size by promoting cell proliferation (Itoh *et al*., 1998; Anastasiou *et al*., 2007; Wang *et al*., 2008; Eriksson *et al*., 2010; Sun *et al*., 2017). At*KLUH* is expressed in the meristem and it was suggested that AtKLUH produces a mobile signal that promotes cell proliferation in distal organs (Anastasiou *et al*., 2007; Poretska *et al*., 2020), however the molecular identity of this signal has not been identified. A single member of subclade A is found in *M. polymorpha*, Mp*CYP78A101* (Mp3g23930). Like Mp*CYP78E1*, this gene is expressed in the thallus notch (Hung *et al*., 2024); however, the authors reported no detectable defective phenotype in loss of function mutants and hypothesise that Mp*CYP78E1* and Mp*CYP78A101* may act redundantly (Hung *et al*., 2024). The enzymatic function of CYP78 family proteins such as MpCYP78E1 therefore remains an open question.

The regulatory networks controlling axillary branching are well understood, but comparatively little is known about the regulation of dichotomous branching. Surgical experiments have demonstrated that an apical auxin source represses meristem activity in *M. polymorpha,* lycophytes and seed plants; a process termed apical dominance (Thimann and Skoog, 1933; Davidonis and Munroe, 1972; Maravolo et al., 1976; Spencer *et al*., 2023; Streubel *et al*., 2023). Reactivation of inactive meristems after removal of the active apical meristem can be suppressed by the application of exogenous auxin to the thallus cut site (Davidonis and Munroe, 1972; Streubel *et al*., 2023), suggesting that apical auxin acts as a mobile signal that inhibits the activity of distant meristems. In addition to auxin signalling, other regulators of meristem activity have recently been identified in *M. polymorpha*. MpSPL1 represses meristem activity and is controlled by the repressor of light signalling MpPHYTOCHROME INTERACTING FACTOR (MpPIF). MpCLE2-MpCLAVATA signalling promotes meristem initiation during dichotomy (Hirakawa *et al*., 2020; Takahashi *et al*., 2021), by reducing the expression of the transcription factor, JINGASA (MpJIN) (Takahashi *et al*., 2023). The spatial regulation and interaction of these multiple signalling pathways remains to be determined.

We have identified MpCYP78E1 as a novel inhibitor of meristem initiation and activity and speculate that MpCYP78E1 produces or modifies a mobile signal in active meristems. This signal may be part of known signalling networks – auxin-mediated or MpSPL1-mediated apical dominance for example – or may act independently to coordinate shoot branching architecture during the life of the plant.

## ACKNOWLEDGEMENTS

We would like to thank the Protein Technologies Facility, the NGS sequencing facility, the BioOptics Facility, and the Plant Sciences Facility at the Vienna BioCenter for their support and advice, as well as the Media Lab, Lab Support and the administrative staff at the VBC. Thank you to Nidhi Mishra, Eva-Sophie Wallner, Hugh Mulvey, Johannes Rötzer, and Zohar Meir for their help and thoughtful discussion. Thank you to Thea Kongsted for critical revision of the manuscript draft. This work was funded by a grant from the Austrian Academy of Sciences to the Gregor Mendel Institute, a European Research Council advanced grant (DENOVO-P, project no. 787613) to L.D. and a European Research Council MSCA grant (DeNovoMeristem, project no. 101148225) to V.S.

## AUTHOR CONTRIBUTIONS

V.S. and L.D. designed the project. V.S. carried out phenotyping experiments and analysis. C.C. harvested samples for RNA Seq and analysed the RNA Seq data set. M.M. carried out spore transformation to generate transgenic plants. K. J. performed PCR and sequencing to verify lines. N.E. propagated plants for experiments, assisted with line verification and maintained stocks. V.S. and L.D. wrote the manuscript. All authors revised the manuscript.

## METHODS

### Plant material and growth conditions

*Marchantia polymorpha* wild type Takaragaike-1 (Tak-1, male) and Takaragaike-2 (Tak-2, female) accessions were used. Plants were grown in walk-in phytotron chambers for 2 weeks in continuous white light (24 hr light/0 hr dark, 45 µmol m^-2^ s^-1^ at 23°C) followed by 6 weeks in simulated shade (16 hr light/8 hr dark, 50 µmol m^-2^ s^-1^ at 20°C), unless stated otherwise. For the first 2 weeks of growth, plants were grown in sterile conditions on plates containing ½ B5 Gamborg’s Media [½ strength B5 Gamborg + 0.5 g/L MES hydrate + 1 % sucrose + 1 % plant agar, pH 5.5] from gemmae. At week 2, plants were transferred to a compost:vermiculite mixture (3:1) in SacO2 microboxes and transferred back to continuous white light (Fig. S1A) or to simulated shade. For Fig. 6, plants were kept on sterile cellulose sheets on media plates in continuous white light for 3 weeks. To induce gametangiophore formation for crossing, wild type Tak-1 and Tak-2 were grown in simulated shade. Mature sporangia were harvested and then dried for 4 weeks before freezing at -70°C.

### Generation of plasmids for transformation

To generate transcriptional (*pro*Mp*CYP89E1:VENUS-NLS)* and translational (*pro*Mp*CYP89E1:g*Mp*CYP78E1-linker-VENUS)* reporter constructs, a standard GreenGate cloning strategy was used (Lampropoulos *et al*., 2013). Using Tak-1 genomic DNA as a template, a 5001 bp promoter sequence (*pro*Mp*CYP78E1*) located directly upstream of the Mp*CYP78E1* start codon was amplified in two pieces using the following primers: Fragment 1= CLON306_pCYP45078E1-F + CLON307_pCYP45078E1-intR; Fragment 2= CLON308_pCYP45078E1-intF + CLON309_pCYP45078E1-R (see Table S1 for sequences). The two PCR fragments were assembled using the NEBuilder® HiFi DNA Assembly Master Mix into the pGGA vector, which was linearized via PCR using the pGG-A-vecR + pGG-B-vecF primers. The genomic Mp*CYP78E1* gene sequence (*g*Mp*CYP78E1*) of 1924bp (excluding the stop codon) was amplified in three pieces to disable the gRNA/Cas9 binding sites via introduction of silent mutations. The gRNA binding sites (5’-3’ gRNA1: ATCCCAGGGGTTGAAGCATGCGG + gRNA2: CATCGCCGTAACCATCGAGTGGG) were mutated (to ATCCCAGGGGTTGAAGCATGCtG + <u>a</u>AT<u>a</u>GCtGT<u>g</u>ACtAT<u>a</u>GA<u>a</u>GGG) to prevent editing of the transgene when introduced into the CRISPR mutant background. The following primers were used for amplification: Fragment 1= CLON310_gCYP45078E1-Fcor + CLON311_gCYP45078E1-fr1-intR; Fragment 2= CLON312_gCYP45078E1-fr2-intF + CLON313_gCYP45078E1-fr2-intR; Fragment 3= CLON314_gCYP45078E1-fr3-intF + CLON315_gCYP45078E1-R (see Table S1 for sequences). The three PCR fragments were assembled using the NEBuilder® HiFi DNA Assembly Master Mix into the pGGC vector, which was linearized via PCR using the pGG-C-vecR + pGG-D-vecF primers. The linker-VENUS module was published previously (Wallner, Dolan and Bergmann, 2023) and the chlorsulfuron plant resistance module was adapted from the OpenPlant toolkit (Sauret-Güeto *et al*., 2020). The modules were assembled into the pGGZ003 destination vector following the standard protocol. The Mp*CYP78E1* entry modules and translational reporter plasmid were cloned by the Protein Technologies Facility at the Vienna Biocenter (https://www.viennabiocenter.org/vbcf/), and the transcriptional reporter plasmid was assembled according to the GreenGate assembly protocol within the lab (Lampropoulos *et al*., 2013). A predicted gene (Mp1g14160) in the 3’->5’ orientation is located immediately upstream of the Mp*CYP78E1* start site. However, both our transcriptome data and the Marpolbase Expression database indicate that this gene is not expressed in meristems or the thallus.

CRISPR-Cas9 plasmids were produced and verified by the Protein Technologies Facility at the Vienna Biocenter. For Mp*CYP78E1* (Mp1g14150), 2 gRNA target sites were selected (5’-3’ gRNA1: ATCCCAGGGGTTGAAGCATG<u>CGG</u> + gRNA2: CATCGCCGTAACCATCGAGT<u>GGG</u>) immediately before the predicted membrane associating domain and within the CYTOCHROME P450 domain (Fig. S6) using the CHOPCHOP web tool with the basic settings (Labun *et al*., 2019). For Mp*MLP-LIKE423* (Mp3g25440), 2 gRNA target sites were selected (5’-3’ gRNA1: TCTCATGAAGCAAGGTATGC<u>AGG</u> + AACTGTGATGACGGGAGGGA<u>TGG</u>), both within the predicted Bet v1/Major latex protein domain. The PAM is underlined in each gRNA. Plasmids were cloned according to the OpenPlant toolkit protocol (Sauret-Güeto *et al*., 2020).

### Generation of transgenic lines

Spores from a Tak-1 x Tak-2 cross were used for the generation of transgenic lines. Spores were transformed with plasmids described above using the protocol adapted from Ishizaki et al., 2008 and described in Wallner and Dolan, 2024. After transformation, sporelings transformed with reporter constructs (*pro*Mp*CYP89E1:VENUS-NLS* and *pro*Mp*CYP89E1:g*Mp*CYP78E1-VENUS)* were plated on ½ B5 Gamborg’s Media containing 0.5 µM chlorsulfuron and 100 mg/L cefotaxime, whilst sporelings transformed with CRISPR constructs were plated on ½ B5 Gamborg’s Media containing 10 mg/l hygromycin and 100 mg/L cefotaxime. Positive transformants were transferred to fresh selective plates, and then gemmae from these plants were transferred to non-selective plates for phenotyping. Lines were screened for consistent expression patterns in gemmae grown for four days in white light. The presence of the transgene in selected lines was further verified using the “Plant sequencing primers” in Table S1.

To generate complementation lines, the *pro*Mp*CYP78E1:g*Mp*CYP78E1-VENUS* construct was transformed into the Mp*cyp78e1^LOF_g2_14m^* mutant background by thallus transformation using the method developed by Kubota *et al*., 2013 and modified by Mulvey & Dolan, 2023. Plants were grown for 2 weeks from gemmae in white light on media plates. A sterile 4 mm diameter cork borer was used to generate 4 thallus discs from the mature thallus of Tak-1 plants, excluding the notches. Discs were cultured for 3 further days on solid media in white light. GV3101 pSoup agrobacterium cultures containing the construct of interest were grown for 2 days in M51C media at 28 °C in the dark with constant agitation. 1 ml of agrobacterium culture was added to 50 ml of M51C media in a 125 ml Erlenmeyer flask, in addition to 100 µM acetosyringone. 40-50 thallus discs were added to the co-culture media and incubated for 3 days at 23 °C in continuous white light (50-60 µmol m^-2^ s^-1^) with constant agitation (130 rpm). Thallus discs were then washed 3 times in sterile water and incubated for 30 minutes in 1 mg/ml cefotaxime. Discs were plated onto media plates containing 0.5 µM chlorsulfuron and 100 mg/L cefotaxime for selection.

### RNA extraction

For qPCR of loss of function and gain of function lines, plants were grown for 2 weeks in continuous white light before harvesting. Four notches from four different plants were harvested per sample, and tissue was frozen immediately in liquid nitrogen before tissue homogenisation with a 5 mm metal bead and a Retsch MM 400 tissue homogeniser. A RNeasy® Plant Mini Kit was used to extract RNA from homogenised samples. The kit protocol was followed, with the following modifications: Buffer RLT was used, and an on-column DNase treatment was performed for 15 minutes in between two buffer RW1 washes. For RNA Seq, plants were grown for 12 days in white light, then transferred to simulated shade for 15 more days. Active and inactive notches were harvested in pairs and RNA extraction was performed on each notch individually. No visible gametangiophores were present at the harvested notches. Inactive meristems were categorised into three stages based on the distance to their neighbouring active notch; stage 1 inactive notches were < 3 mm from the active notch, stage 2 notches were between 3-5 mm from the active notch, and stage 3 notches were > 5 mm from the active notch. Meristems were harvested from the 3^rd^ plastochron and were immediately frozen in liquid nitrogen.

### cDNA synthesis and qPCR

To quantify gene expression in LOF and GOF samples, cDNA synthesis was first performed using the LunaScript® RT SuperMix Kit according to the manufacturer’s protocol. qPCR was performed using the Luna® Universal qPCR Master Mix and a LightCycler® 96 machine. 2 technical replicates and 3 biological replicates were performed for each line. *ADENINE PHOSPHORIBOSYLTRANSFERASE* (Mp*APT*; Mp3g25140) was used as a control gene (Saint-Marcoux *et al*., 2015). qPCR primers are stated in Table S1. The Mp*CYP78E1* reverse qPCR primer spans an exon-exon boundary.

### RNA Sequencing and analysis

RNA samples were sent for sequencing to the VBCF (Vienna BioCenter Core Facilities) Next Generation Sequencing (NGS) Facility (https://www.viennabiocenter.org/vbcf/). Library preparation, quality control and sequencing were performed by the facility. All samples passed quality checks as determined by the facility (RIN > 7 and DV200 > 70 %). Poly-A mRNA was isolated with the NEBNext® Poly(A) mRNA Magnetic Isolation Module (NEB, #E7490) from 500 ng of total RNA. Enriched poly-A mRNA sequencing libraries were prepared using the NEBNext® Ultra™ II Directional RNA Library Prep Kit for Illumina® following the kit instructions. The fragment size of the libraries was assessed using the NGS HS analysis kit on an Agilent fragment analyzer system. The library concentrations were quantified with a Roche KAPA Kit by qPCR.

RNA libraries were sequenced on an Illumina NovaSeq 6000 S1 flowcell using 50 bp paired-end reads. The following analyses were performed using the CLIP high-performance computing cluster at the Vienna BioCenter. Raw reads were trimmed to remove Illumina adaptors and low-quality reads using Trimmomatic v0.38 (Bolger *et al*., 2014). Read counts were quantified using Salmon v1.5.2 (Patro *et al*., 2017) in quasi-mapping-based mode against a salmon transcriptome index built from the MpTak_v6.1 transcriptome.

All subsequent analyses were performed using R v4.2.0. Transcript-level Salmon quantification files were imported and gene-level summarization was performed using tximport v1.26.1 (Soneson *et al*., 2015). Gene-level TPM values (transcripts per million) were extracted with tximport using a transcript to gene (tx2gene) mapping file constructed from the MpTak_v6.1 transcriptome. Gene-level TPMs for the Mp1g14150 gene were transformed as log2(TPM + 1) in Fig. 2C to visualize gene expression across samples.

To identify genes that were differentially expressed between inactive and active meristems at each of the three stages, differential gene expression analysis was carried out using DESeq2 v1.38.3 (Love *et al*., 2014). The design formula “ ∼ pair + activity ” was used to identify genes differentially expressed between inactive and active meristems while controlling for variability between sample pairs. Significantly differentially expressed genes were defined to have a Log2 Fold Change < -2 or > 2, and a p_adj_ < 0.05.

### Phylogenetic analysis

The Mp1g14150 protein sequence was used in SHOOT.bio to generate a phylogenetic tree of this orthogroup + 2 levels (Emms and Kelly, 2022). The tree was generated using 30 plant species plus 3 red algal outgroups, as specified by the SHOOT.bio plant subset.

### Tissue fixation and clearing

Samples were fixed and cleared following a modified version of the protocol described by Mulvey & Dolan, 2023. Notches were harvested and transferred immediately to 10 % formalin solution (10 % neutral buffered, 4 % w/v formaldehyde) + 0.1 % Brij® L23 solution. Samples were vacuum infiltrated for 25 minutes, followed by another 25 minutes (1 hour total). Samples were then washed three times with phosphate buffered saline (PBS), before incubation in Clear-See α solution [10 % (w/v) xylitol + 15 % (w/v) sodium deoxycholate + 25 % (w/v) urea + 50 mM sodium sulfite anhydrous]. Samples were vacuum infiltrated for 25 mins, before incubation in the dark overnight. The next day, the Clear-See α solution was replaced. Samples were stored in the dark on a rocker until imaging (minimum 5 days).

### Imaging, microscopy and image analysis

Live plants on soil were imaged with a Canon EOS 80D camera with a Canon EF 50mm Lens, and live plants on media were imaged with the Keyence VHX-7000 digital microscope with the VH-ZST and VH-Z00R/W/T objectives. The 3D tilescan function was performed on the Keyence microscope for images in Fig. 6. Fixed and cleared samples were mounted following the protocol described in Spencer *et al*., 2024. The day before imaging, the cell wall stain SR2200 was added directly to the samples in clear-see solution to a final concentration of 0.2 %. Samples were mounted on glass slides in 0.25 mm thick gene frames filled with Clear-See solution and sealed with a #1.5 coverslip. Images were acquired with an inverted point scanning Zeiss LSM880 confocal microscope using the x10/0.45 plan-apochromat objective and MBS 458/514 and MBS -405 filters. To detect SR2200 stain, a 405 nm diode laser (1 % for all sample) was used for excitation and signal was detected between 419-499 nm with a 25 µm pinhole (∼1 AU). To detect VENUS signal, a 514 nm argon laser (8 % for transcriptional reporter line _8f, 18 % for transcriptional line _1m, and 20 % for translational reporter lines) was used for excitation and VENUS signal was detected between 526-579nm with a 31 µm pinhole (∼1 AU). The two channels were scanned sequentially in all imaging and the track was switched after each z-stack. Z -stacks of 50 slices with a 2.4 µm interval were acquired for VENUS lines. All image analysis was performed using Fiji. Z-projections were produced for the VENUS channel using the Z-projection: Maximum Intensity function in Fiji.

### Phenotyping, statistical analysis and data presentation

Notches were counted as a proxy for meristem number. To quantify the numbers of active and inactive meristems of mature plants, meristems were counted as active when they were positioned at the thallus apex, and inactive when positioned beneath the thallus apex, on a side branch that was less than 10 mm long. When quantifying meristem number in Fig. 5, samples were excluded if they were growing into the agar and notches were not visible. The number of notches was counted on the largest half of the thallus body. Any notches on lobes that were growing out of the central thallus body were excluded from the analysis. All figures were made in Excel and Inkscape and statistical analysis was performed using R-studio. For Fig. 6, the grey background was removed in Microsoft Powerpoint before the image was transferred to Inkscape, and the original images are displayed in Fig. S9. A one-way ANOVA with Tukey’s HSD multiple comparison test was performed when data were normally distributed with equal variance. When these assumptions were not met, a Kruskal Wallis test with Dunn’s multiple comparison was performed. Experiments were carried out once unless stated otherwise.

### Reagents, equipment and software

See Table S1 for details.

## Data and code availability

The RNA Seq raw data is available on the NCBI GEO repository (accession: GSE295341). The code for the RNA Seq analysis is available at https://github.com/calcasey/m_polymorpha_meristem_activity_transcriptomics.

---

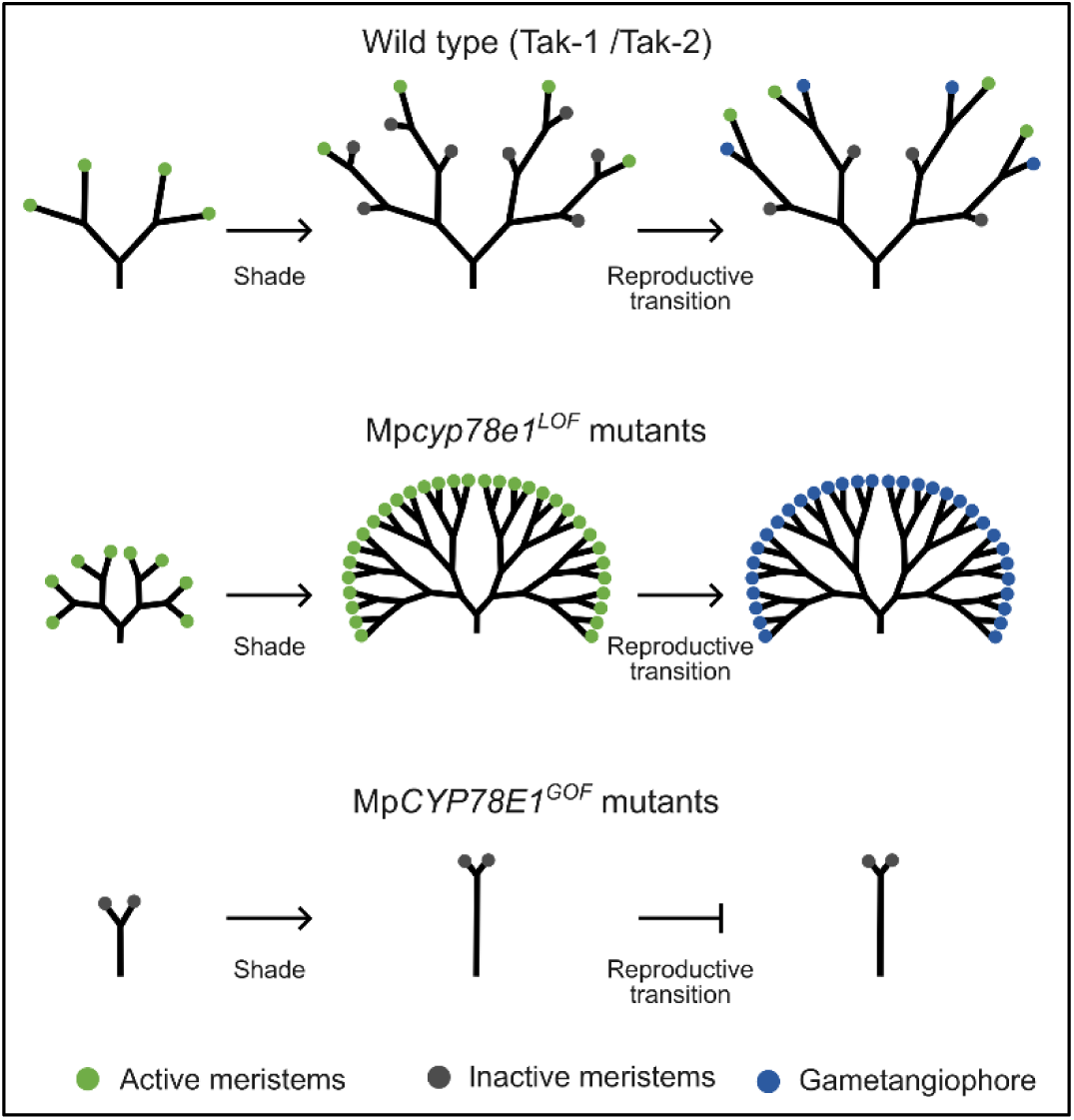
Summary diagram: Schematic to summarise the architectures of wild type, Mp*cyp78E1^LOF^* lines and Mp*CYP78E1^GOF^*lines. The meristems of wild type plants are all active when grown in white light. After transfer to simulated shade, both active and inactive meristems are formed, and only the most apical active meristem produces gametophores during the reproductive transition. The neighbouring meristem activates and replaces the leading growth axis. In Mp*cyp78e1^LOF^* lines, more meristems are produced in white light due to increased meristem initiation by dichotomy. After transfer to shade, all meristems grow equally and are unable to inactive. During the reproductive transition, all meristems transition to gametophores, supporting the hypothesis that all meristems are active in these mutants. Mp*CYP78E1^GOF^* lines grow more slowly and undergo fewer dichotomies in white light. After transfer to simulated shade, no reproductive structures are formed, supporting the hypothesis that MpCYP78E1 is an inhibitor of meristem activity as meristems are inhibited in the Mp*CYP78E1^GOF^* lines. Green circles= active meristems. Dark grey circles= inactive meristems. Blue circles= gametangiophores.

**Figure S1:**
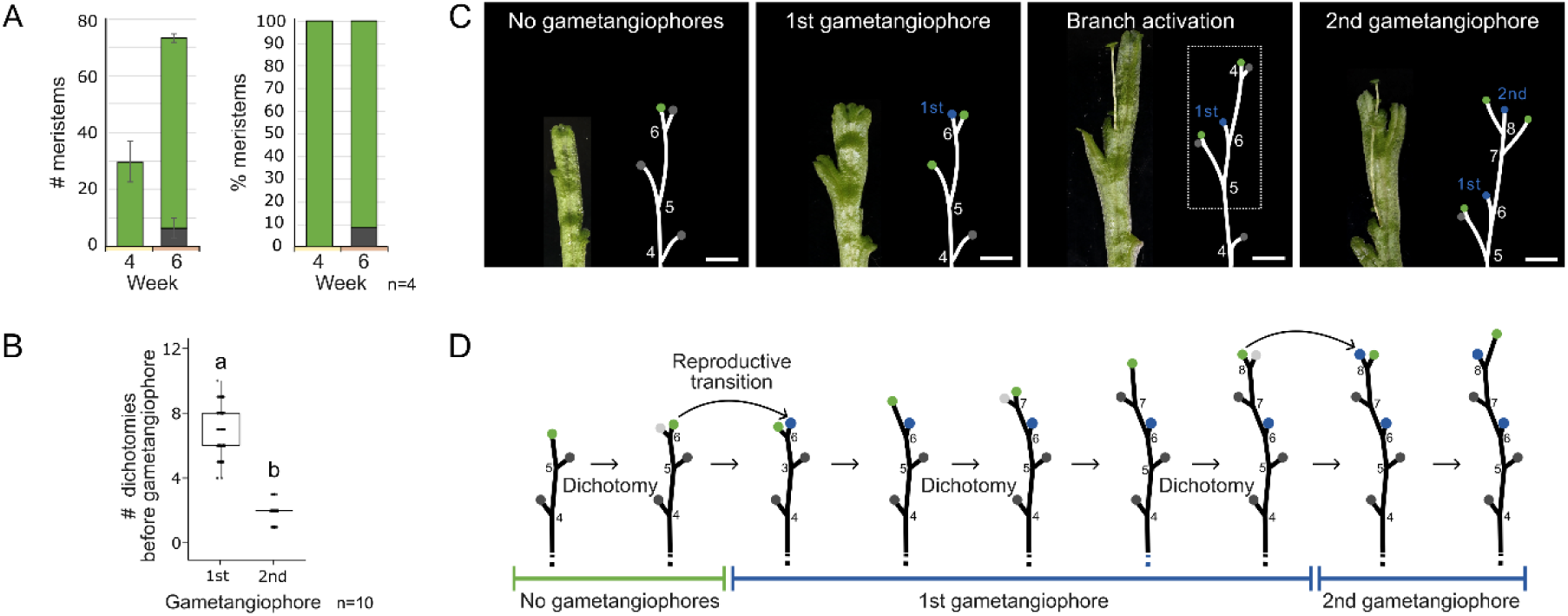
Patterns of wild type Tak-1 gametangiophore production in simulated shade. (**A**) Quantification of the number of active meristems (green), inactive meristems (dark grey), and gametangiophores (blue) after 4 and 6 weeks of growth in white light. All meristems remain active at week 4. Inactive meristems are formed at week 6 as the plant ages. (**B**) Quantification of the total number of dichotomies between gametangiophores in wild type Tak-1 plants grown for 2 weeks in white light followed by 6 weeks in simulated shade. The first gametangiophore is produced after an average of 6.7 dichotomies, whilst the 2^nd^ gametangiophore of the same branch is produced after an average of 1.9 more dichotomies. A Kruskal-Wallis test with Dunn’s multiple comparison was performed [χ^2^(1)=112.0, p=3.33E-26]. n= 104 for the 1^st^ gametangiophore and n=56 for the 2^nd^ gametangiophore, totalled from 10 plants. Box and whisker plots represent the median and the lower and upper quartiles. (**C**) The position of gametangiophores was recorded at week 8. Based on the predictable alternation of inactive meristems to the left and right, the gametangiophores are predicted to form from the active meristem. (**D**) Schematic of the position of gametangiophore production. Only the active meristem produces a gametangiophore. The second gametangiophore is formed after two more dichotomy events. Green circles= active meristems. Light grey circles= future inactive meristems. Dark grey circles= inactive meristems. Blue circles= gametangiophores. Scale bars = 10 mm.

**Figure S2:**
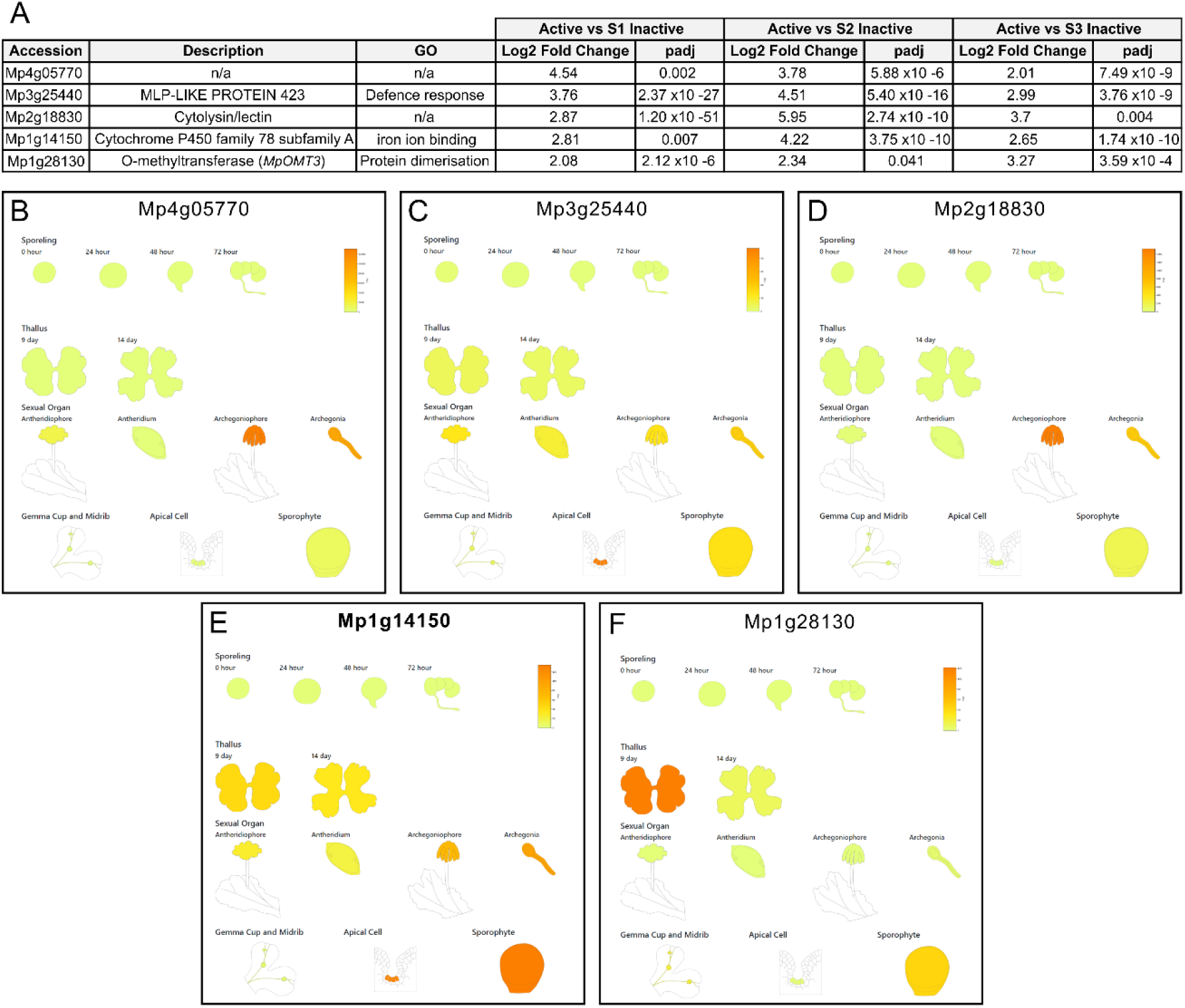
RNA Seq data and MarpolBase expression atlas data for active meristem enriched genes. (**A**) Table showing Log2 Fold Change and padj values for the top 5 genes that were enriched in the active meristem compared to the inactive meristem at all three stages of inactivation. (**B**-**F**) The MarpolBase expression atlas shows that Mp3g25440 (C) and Mp1g14150 (E) were enriched in the meristem stem cell niche compared to other tissues. For this reason, these genes were chosen for further analysis and CRISPR loss of function mutants were made.

**Figure S3:**
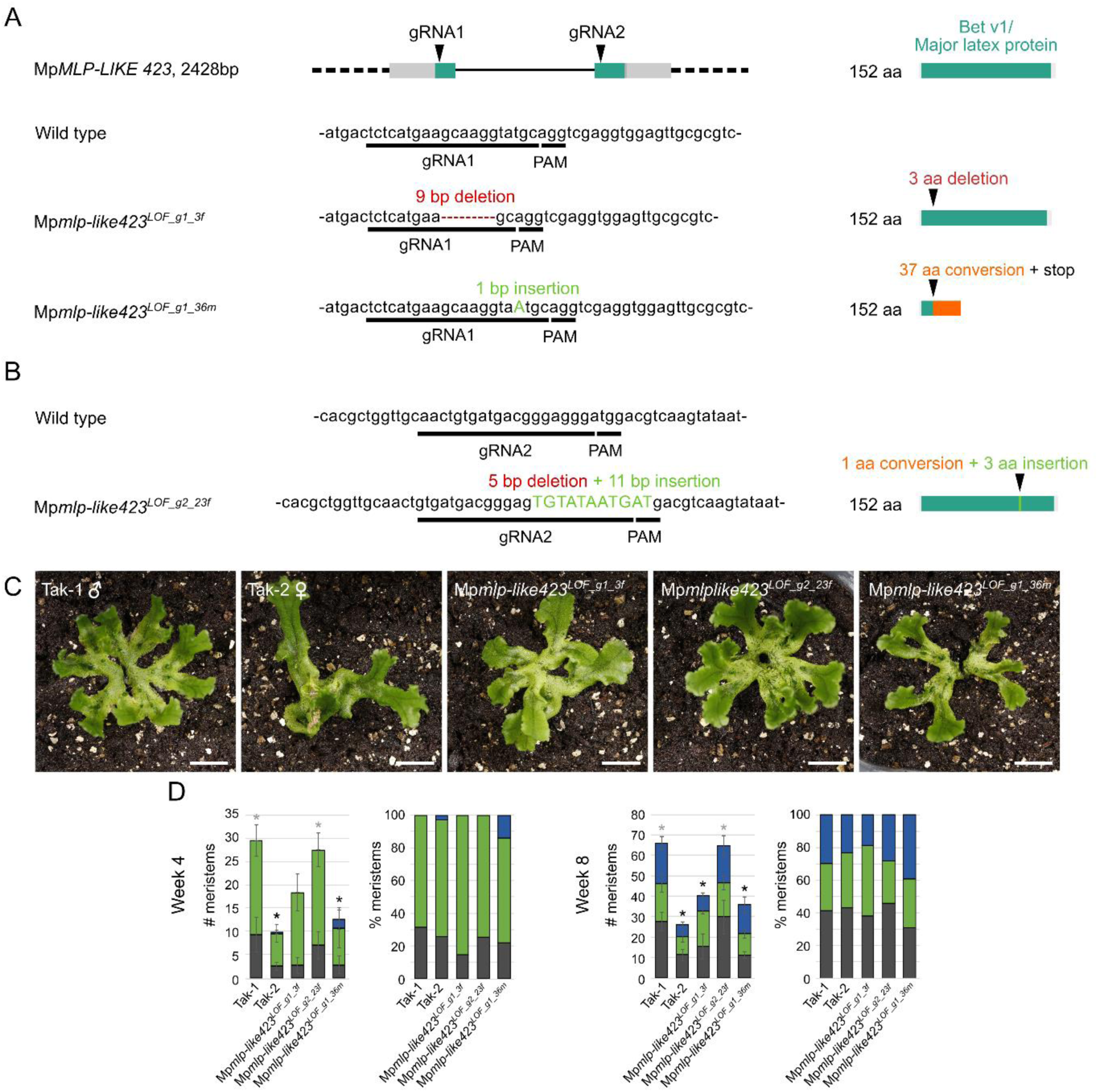
Mp*mlp-like423^lof^* (Mp3g25440) CRISPR mutants have no obvious defective phenotype in shade. (**A**-**B**) Mp*MAJOR-LATEX-LIKE423* was identified as a potential meristem activity regulator in the RNA Seq dataset (Fig. S2). To generate loss of function mutants, gRNAs (gRNA1 + gRNA2) were designed within the predicted Major Latex Protein domain (teal). The DNA sequences and predicted amino acid sequences of wild type Tak-1 plants and the 3 independent mutants are shown. (**C**) Photographs of wild type Tak-1 and Tak-2 and 3 independent lines grown for 2 weeks in white light followed by 2 weeks in simulated shade. Wild type photos are also used in Fig. 4 as these experiments were performed simultaneously. Scale bars = 10 mm. (**D**) At week 4 and 8, the total number of meristems and the percentage of each meristem type (active= green, inactive= dark grey, reproductive= blue) in Mp*mlp-like423^LOF^*mutants was intermediate of wild type Tak-1 and Tak-2 plants. One-way ANOVA with Tukey’s HSD multiple comparison tests were performed for the total number of meristems at week 4 and 8 respectively (F(4,14)=8.823, p=8.97E-4; F(4,13)=14.938, p=8.72E-5). n= 4 for all lines except n= 3 for 121_3f (week 4 and 8) and 121_36m (week 8). Error bars indicate standard deviation. Black * denotes a significant difference to Tak-1 (p-value <0.05) and grey * denotes a significant difference to Tak-2 (p-value <0.05).

**Figure S4:**
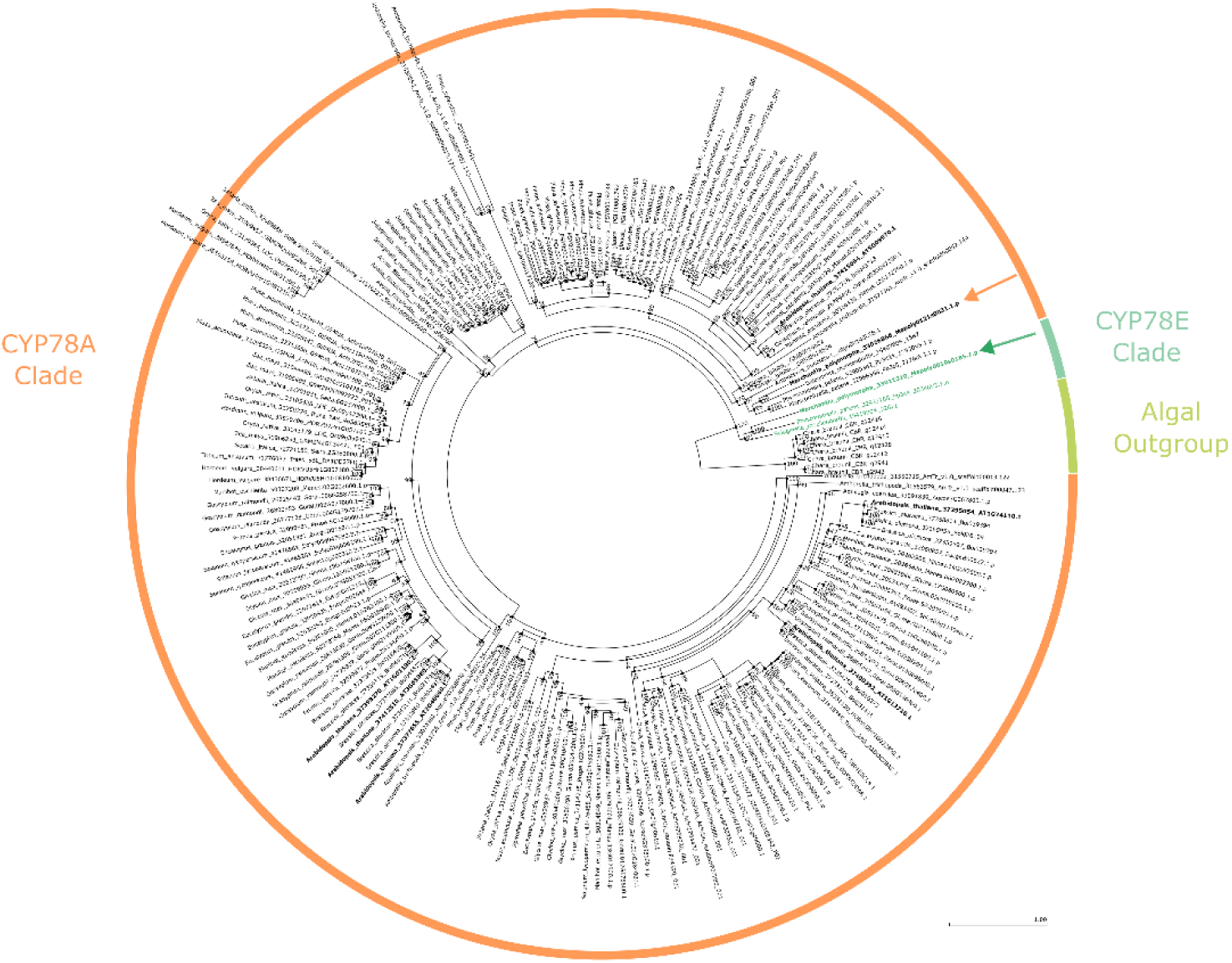
Cytochrome P450 Family 78 phylogeny. Phylogenetic tree of CYTOCHROME P450 family 78 generated by SHOOT.bio using OrthoFinder. Mp*CYP78E1* (green arrow) is within the CYP78 E subfamily. The closest sister group is the CYP78 A subfamily, which includes the *M. polymorpha* homolog, Mp*CYP78A101* (Mp3g23930, orange arrow), as well as *Arabidopsis thaliana* genes such as At*KLUH*.

**Figure S5:**
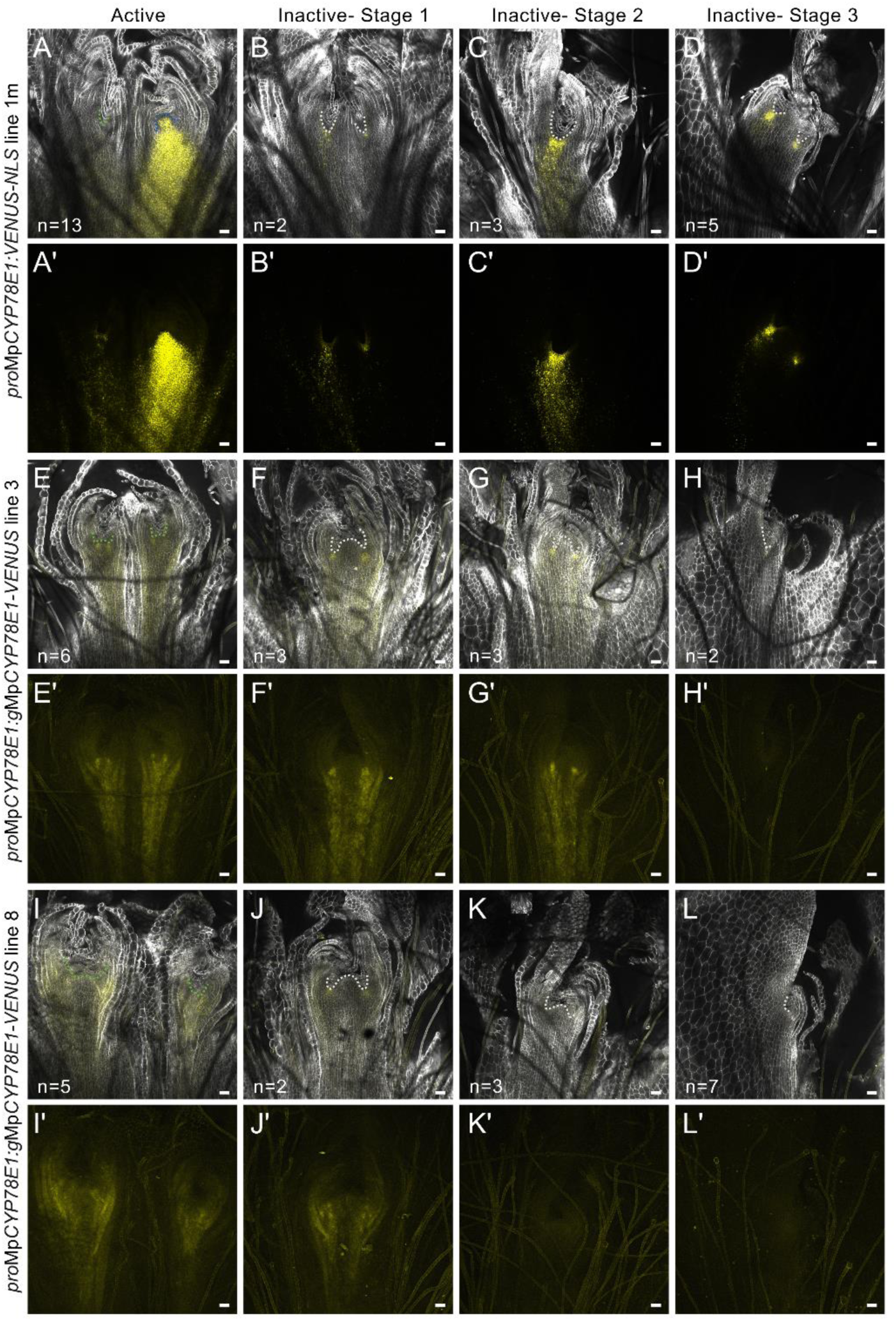
2^nd^ Independent line of Mp*CYP78E1* showing consistent results with Figure 3. (**A**-**D’**) Confocal images of fixed and cleared meristems of *pro*Mp*CYP78E1:VENUS-NLS* line_1m at different stages of meristem inactivation. A, B, C, and D show optical sections in the frontal plane. A’, B’, C’ and D’ show z-projections of the reporter signal across a Z-stack of the meristem. There is strong signal accumulation around each stem cell niche and this extends basally from the meristem. Signal is enriched in the gametangiophore meristem. The experiment was repeated twice and showed consistent results. Sample number is pooled from both replicates and is shown in the figure. (**E**-**L**) Confocal images of fixed and cleared meristems of *pro*Mp*CYP78E1:*g*MpCYP78E1-VENUS* line 3 (E-H’) and line 8 (I-L’) at different stages of meristem inactivation. E, F, G, H, I, J, K and L show optical sections in the frontal plane. E’, F’, G’, H’, I’, J’, K’ and L’ show z-projections of the reporter signal across a Z-stack of the meristem. Cell wall = white. VENUS signal = yellow. Scale bars = 50 mm.

**Figure S6:**
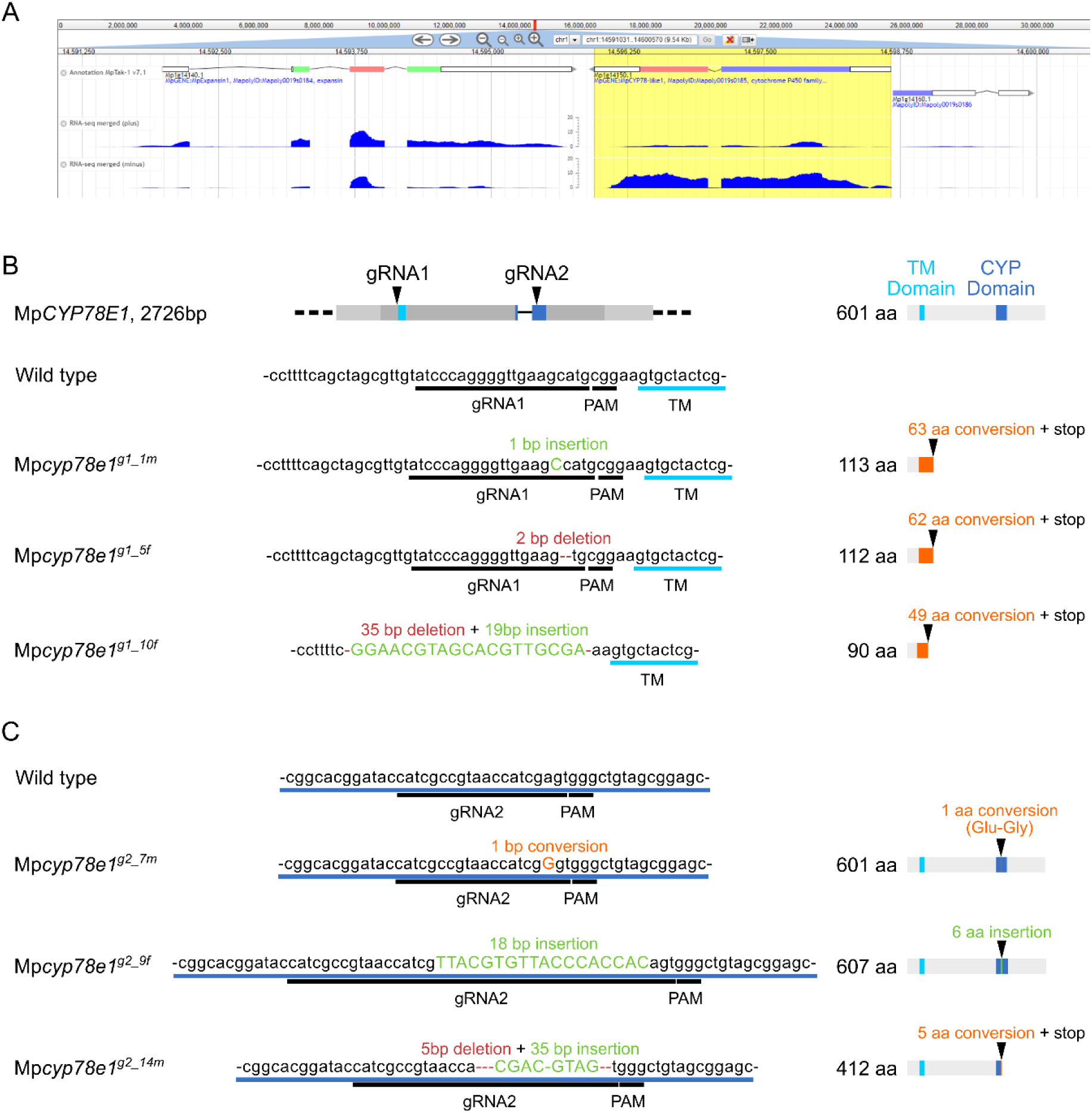
Mp*CYP78E1^LOF^* CRISPR mutant description. (**A**) Genomic location of Mp1g14150 as shown in the Marchantia.info browser. (**B, C**) To generate loss of function mutants, gRNAs (gRNA1 + gRNA2) were designed to target just before the transmembrane domain (TM, pale blue) (B) and inside the CYP domain (CYP, dark blue) (C) respectively. 3 independent mutant lines were selected for each gRNA. The DNA sequences and predicted amino acid sequences of wild type Tak-1 plants and the 6 independent mutants are shown.

**Figure S7:**
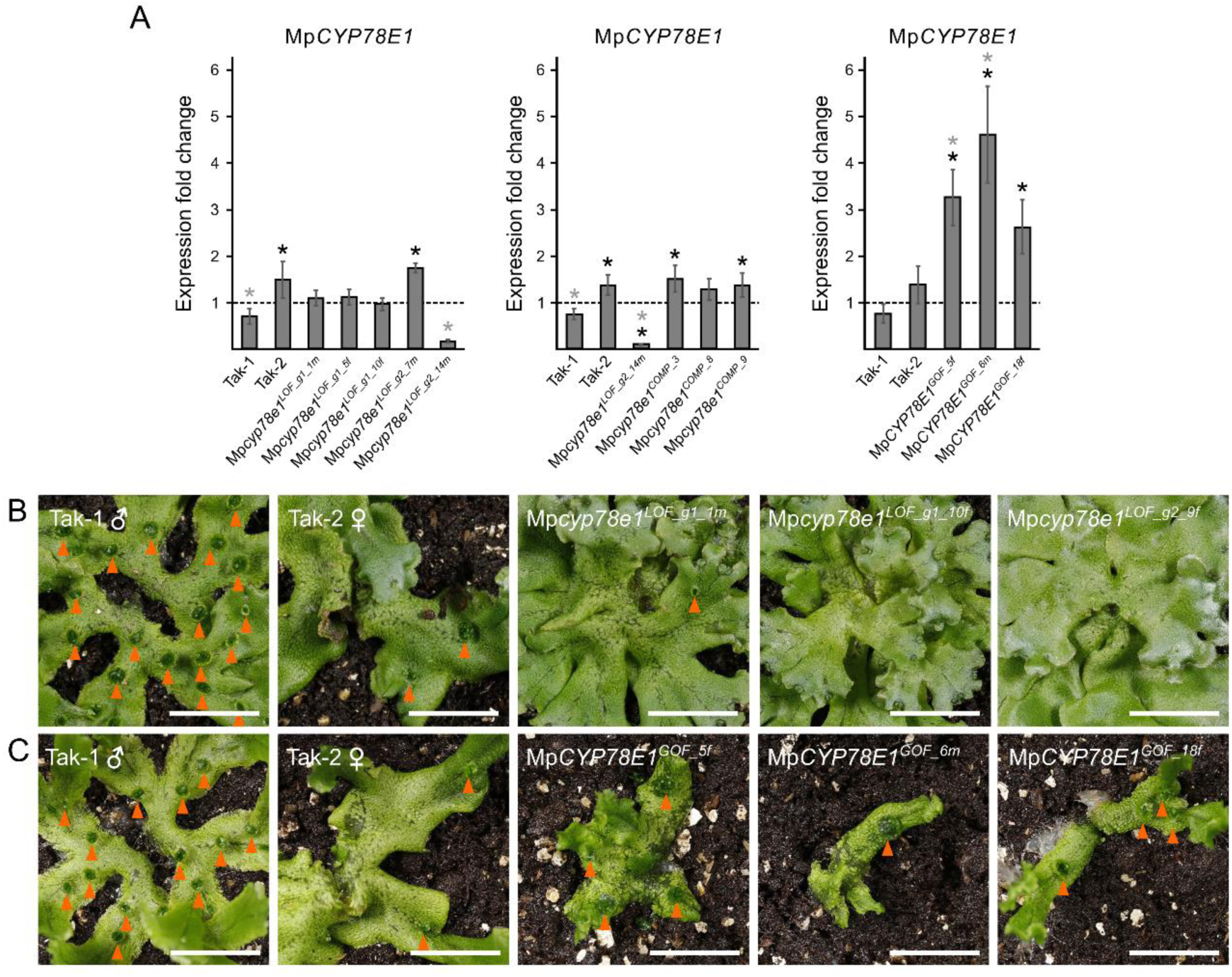
qPCR and gemma cup phenotypes of Mp*CYP78E1^LOF^* and Mp*CYP78E1^GOF^* mutants. (**A**) Quantification of Mp*CYP78E1* expression in loss of function (LOF), gain of function (GOF), and complementation (COMP) lines by qPCR. One-way ANOVA with Tukey’s HSD multiple comparison tests were performed [F(6,14)=21.169, p=2.85E-06; F(5,12)=20.69, p=1.61E-05; F(4,10)=17.632, p=1.59E-04]. 2 technical replicates and 3 biological replicates were carried out per transgenic line and error bars indicate standard deviation. Black * denotes a significant difference to Tak-1 (p-value <0.05) and grey * denotes a significant difference to Tak-2 (p-value <0.05). (**B**) The gemma cups (orange arrowheads) of Mp*cyp78e1^LOF^* mutants appeared smaller than wild type Tak-1 and Tak-2. (**C**) The gemma cups of Mp*CYP78E1^GOF^*mutants were larger than wild type. Scale bars = 10 mm.

**Figure S8:**
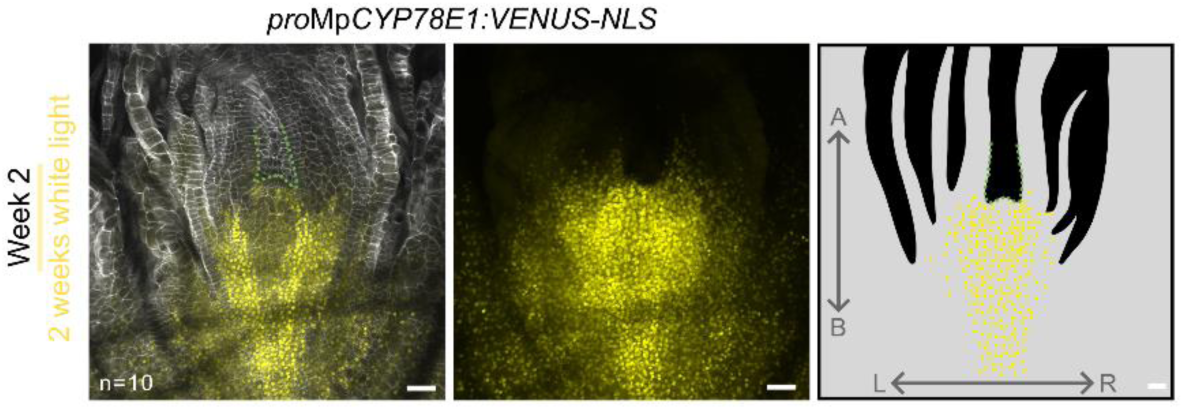
2^nd^ Independent line of *pro*Mp*CYP78E1:VENUS-NLS* showing consistent results with. **Figure 5** Confocal image of fixed and cleared meristem of *pro*Mp*CYP78E1:VENUS-NLS* line_1m grown for 2 weeks in white light. Left, an optical section in the frontal plane; right, a z-projection of the reporter signal across a Z-stack of the meristem. There is strong signal accumulation around each meristem and extending back from the meristem. Cell wall= white. *pro*Mp*CYP78E1:VENUS-NLS* signal= yellow. Scale bars = 50 µm.

**Figure S9:**
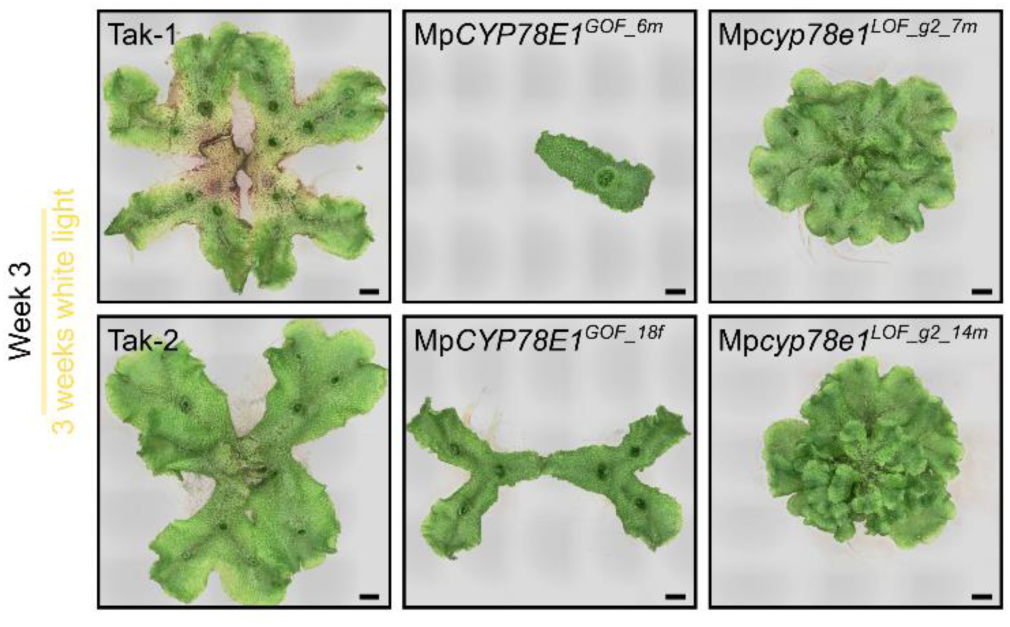
Images of wild type, Mp*CYP78E1^GOF^* lines and Mp*cyp78e1^LOF^* lines shown in Figure 6. (**A**) Photographs of wild type Tak-1, Tak-2, Mp*CYP78E1^GOF^* lines and Mp*cyp78e1^LOF^* lines after three weeks of growth in white light. These are the originals of the images shown in figure 6, but without the background removed. Scale bars = 2.5 mm.

**Table S1:**
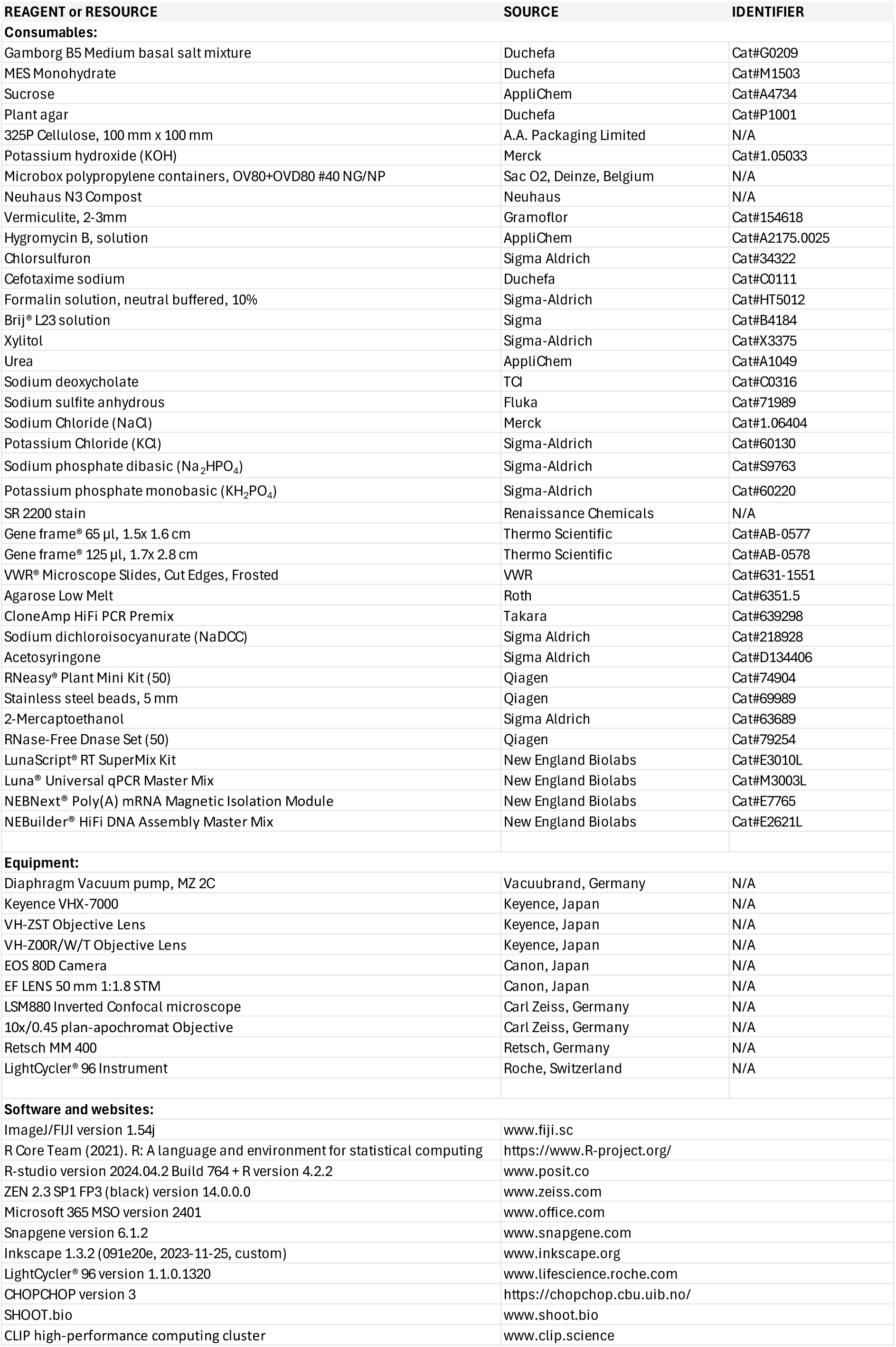
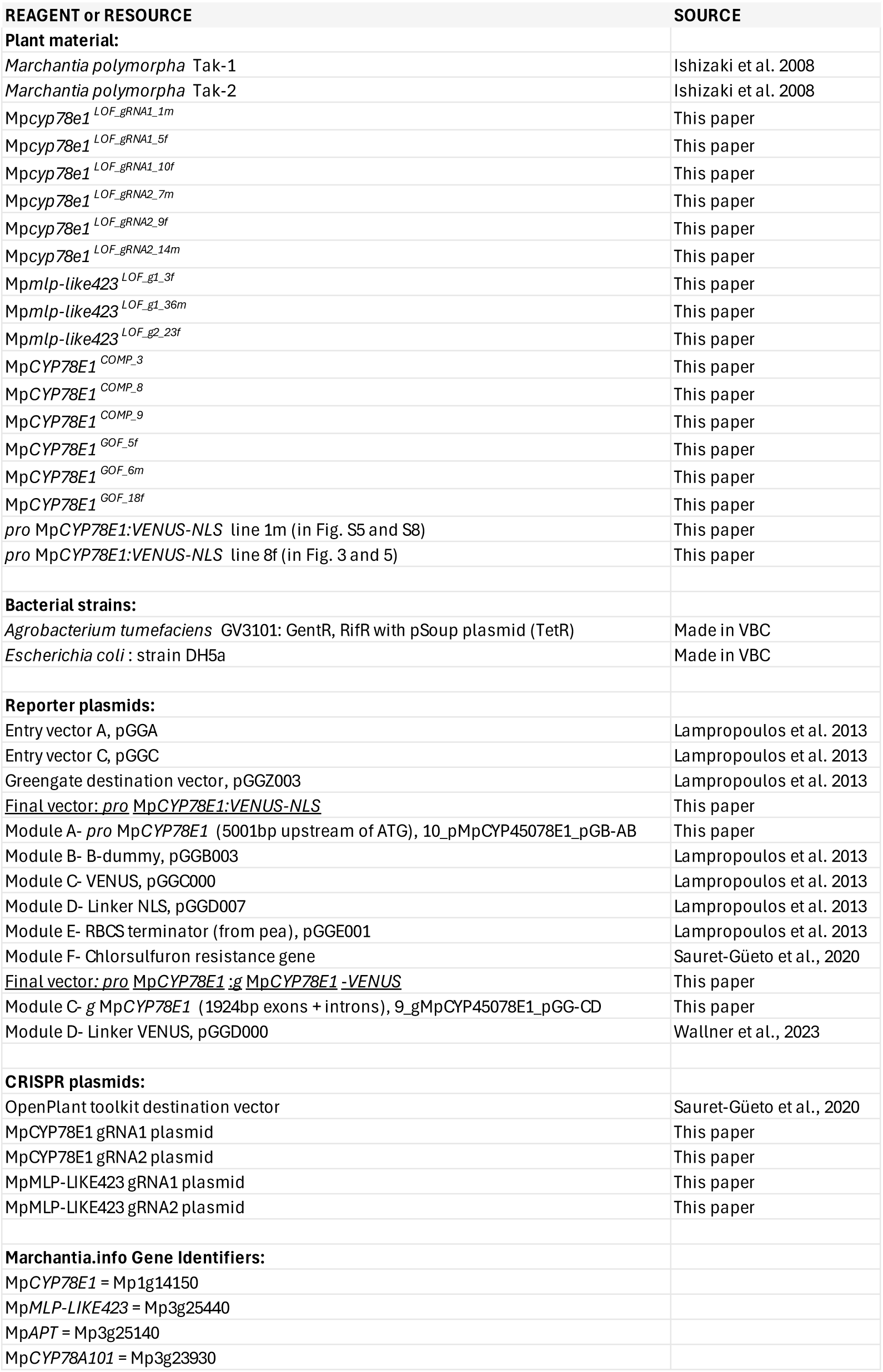
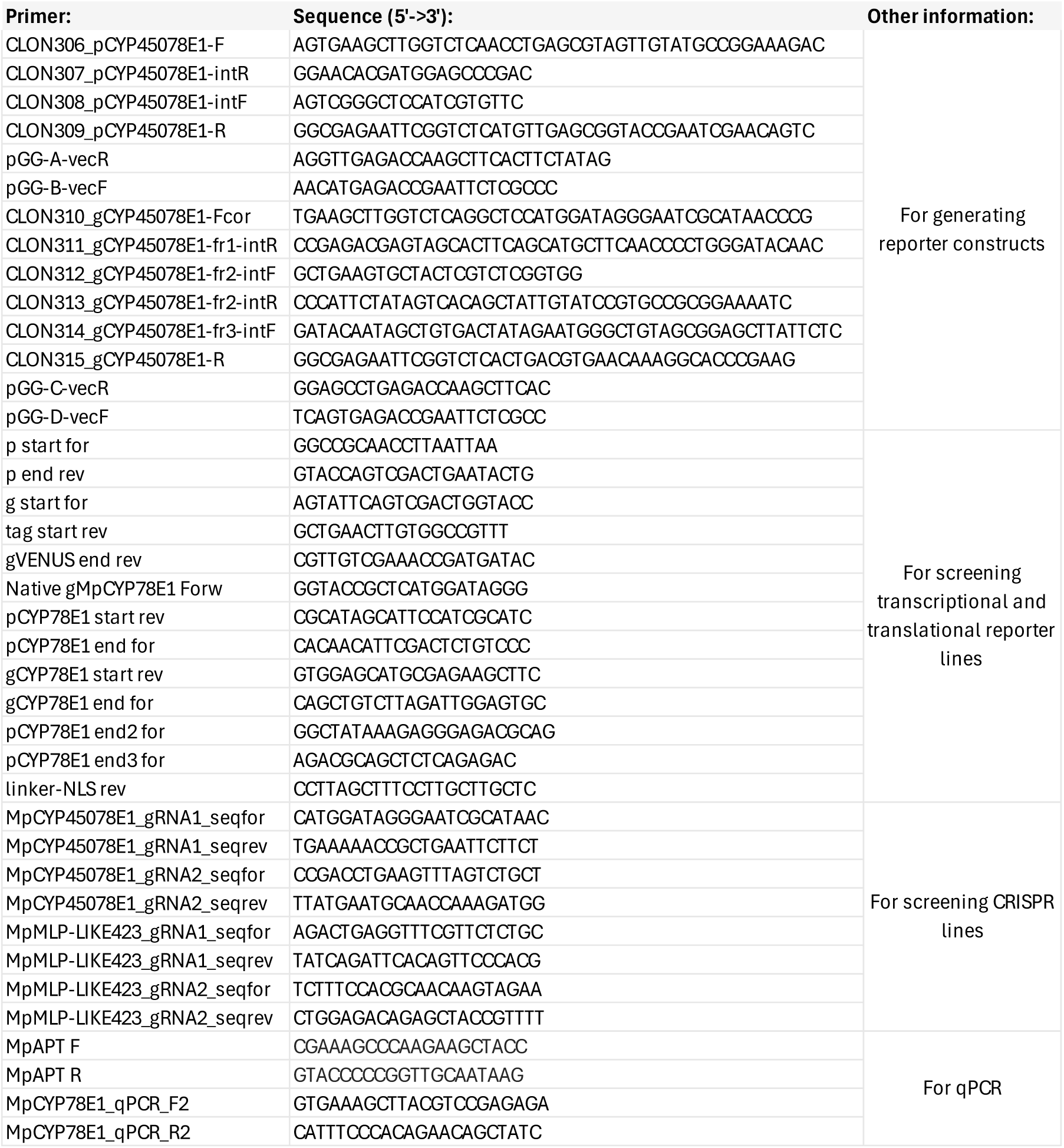
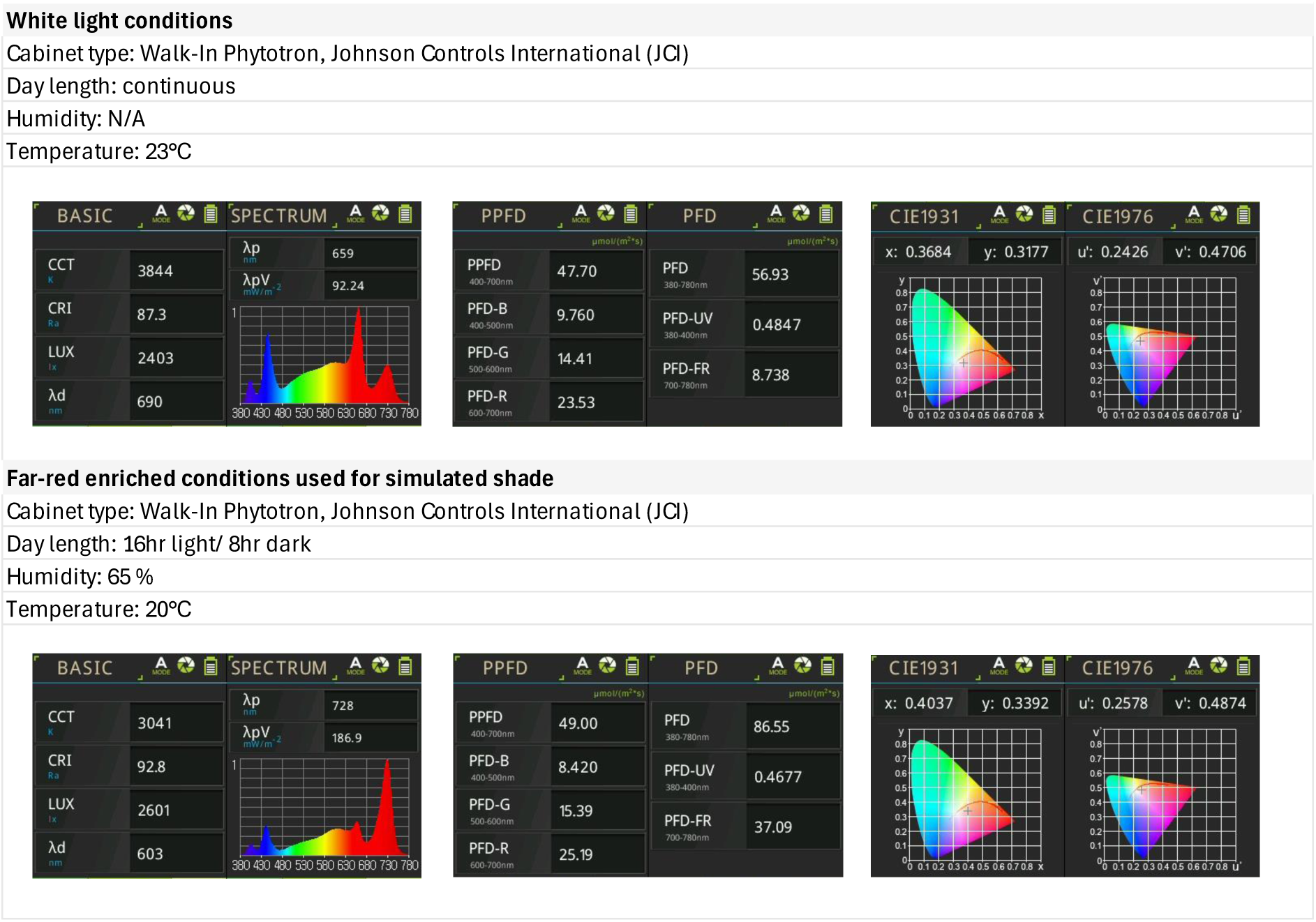

